# From Value to Saliency: Neural Computations of Subjective Value under Uncertainty in PTSD

**DOI:** 10.1101/2020.04.14.041467

**Authors:** Ruonan Jia, Lital Ruderman, Charles Gordon, Daniel Ehrlich, Mark Horvath, Serena Mirchandani, Clara DeFontes, Steven Southwick, John H. Krystal, Ilan Harpaz-Rotem, Ifat Levy

**Author notes:** These authors contributed equally (Ilan Harpaz-Rotem, Ifat Levy). Corresponding authors: R.J.; I.L.

## Abstract

Military personnel engaged in combat are vulnerable to Posttraumatic Stress Disorder (PTSD), following traumatic experiences in the battlefield. Prior research has mostly employed fear-related paradigms to unravel neural underpinnings of fear dysregulation in individuals with PTSD. The ability to acquire and update fear responses depends critically on the individual’s ability to cope with uncertainty, yet the role of individual uncertainty attitudes in the development of trauma-related psychopathology has hardly been examined. Here, we investigated the association between PTSD-related alterations and the subjective valuation of uncertain outcomes during decision-making. We used a monetary gambling paradigm inspired by behavioral economics in conjunction with fMRI and explored neural markers of both vulnerability and resilience to PTSD in a group of combat veterans. Behaviorally, PTSD symptom severity was associated with increased aversion to uncertainty. Neurally, activity in the ventromedial prefrontal cortex (vmPFC) during valuation of uncertain options was associated with PTSD symptoms, an effect which was specifically driven by numbing symptoms. Moreover, the neural encoding of the subjective value of those uncertain options was markedly different in the brains of veterans diagnosed with PTSD, compared to veterans who experienced trauma but did not develop PTSD. Most notably, veterans with PTSD exhibited enhanced representations of the saliency of rewards and punishments in the neural valuation system, especially in ventral striatum, compared with trauma-exposed controls. Our results point to a link between the function of the valuation system under uncertainty and the development and maintenance of PTSD symptoms, and stress the significance of studying reward processes in PTSD.

## Introduction

Following a life-threatening experience, some individuals develop Posttraumatic Stress Disorder (PTSD) symptoms, which can be emotionally, socially and vocationally disabling. These symptoms include re-experiencing the traumatic event, avoidance of trauma reminders, and exaggerated arousal and reactivity, as well as emotional numbing (losing interest in significant activities, having difficulty experiencing happiness or love, and feeling distant from others (1)). While medications and psychotherapy help some individuals, many people with PTSD remain symptomatic following treatment (2). A better understanding of the neural basis of PTSD is crucial, as it can inform new approaches to individualized treatment.

Soldiers in combat face highly uncertain life-threatening events, which are uncontrollable (3), and that may result in serious injury to themselves or death of teammates. An individual’s attitude towards uncertainty and their capacity to handle uncertainty may therefore affect one’s ability to cope with potentially traumatic events. The notion of uncertainty was incorporated in studies of fear-learning attempting to unravel the behavioral and neural mechanisms of PTSD (4,5). Participants in these studies encountered probabilistic deliveries of adverse outcomes (e.g. electric shocks), and their ability to predict these outcomes was measured (e.g. by their skin conductance responses). In a separate line of work, using a behavioral economic framework, our group showed increased aversion to ambiguity (an uncertain situation where outcome probabilities are not known) in combat veterans with PTSD, compared to trauma-exposed veterans without PTSD, when choosing between potential monetary losses (6). This aversion to uncertainty demonstrated in situations unrelated to the trauma, may also contribute to the exaggerated behavior in fear conditioning paradigms, and to the development and maintenance of PTSD symptoms. Further understanding of this aversion to uncertainty could provide evidence for targeting uncertainty in behavioral interventions, to improve the daily decision-making and well-being of individuals with PTSD.

One possibility is that the increased aversion to uncertainty reflects alterations in the neural computations of subjective value in the brains of individuals who developed PTSD following trauma exposure. In the general population, a network of brain regions was implicated in valuation and decision making, including the ventromedial prefrontal cortex (vmPFC), anterior cingulate cortex (ACC), posterior cingulate cortex (PCC), dorsolateral prefrontal cortex (dlPFC), ventral striatum, amygdala, and thalamus (7,8). Extensive evidence suggests that the subjective value of rewards is encoded in this network (9–11), and there is also some (12–15), although less conclusive (16–20), evidence, for encoding of subjective value of punishments in the same areas. Although there is some evidence for changes in the neural processing of monetary outcomes in individuals with PTSD (21,22), we do not know how uncertain decision values are encoded in the brains of these individuals. Moreover, as far as we know, the neural encoding of ambiguous losses (as opposed to gains) has not been investigated even in the general population.

Here we combined a simple economic task with functional MRI and computational modeling to examine the neural encoding of subjective value under risk and ambiguity, and the alterations in this encoding in individuals exposed to trauma. We compared combat veterans with PTSD to those who did not develop PTSD symptoms (trauma-exposed controls), and were thus able to investigate both the psychopathology of PTSD and the resilience to PTSD. We find that veterans with PTSD encode the subjective values of uncertain monetary gains and losses in a U-shape manner, with increased activation for both increased gains and increased losses (compatible with saliency encoding). Conversely, trauma-exposed controls encode the same type of subjective values monotonically, with increased activation for increased gains, and decreased activation for increased losses (compatible with value encoding). Our results suggest that this shift from value-encoding to saliency-encoding, especially of ambiguous monetary losses, could be a neural marker for PTSD symptom severity.

## Results

In an fMRI experiment, combat veterans with current PTSD diagnosis and those who never developed PTSD completed a gambling task under four decision conditions on two separate days. Participants chose between a sure monetary outcome (either gaining or losing $5) and an uncertain outcome (either risky or ambiguous gain or loss) (Fig 1B). Participants made decisions about gains and losses in separate blocks in two scanning sessions (Fig 1A). We estimated the attitudes toward risk and ambiguity of each participant through a behavioral model (see Methods) and aimed to understand the influence of PTSD symptom severity on both the behavioral attitudes and the neural mechanisms of valuation.

**Figure 1.**
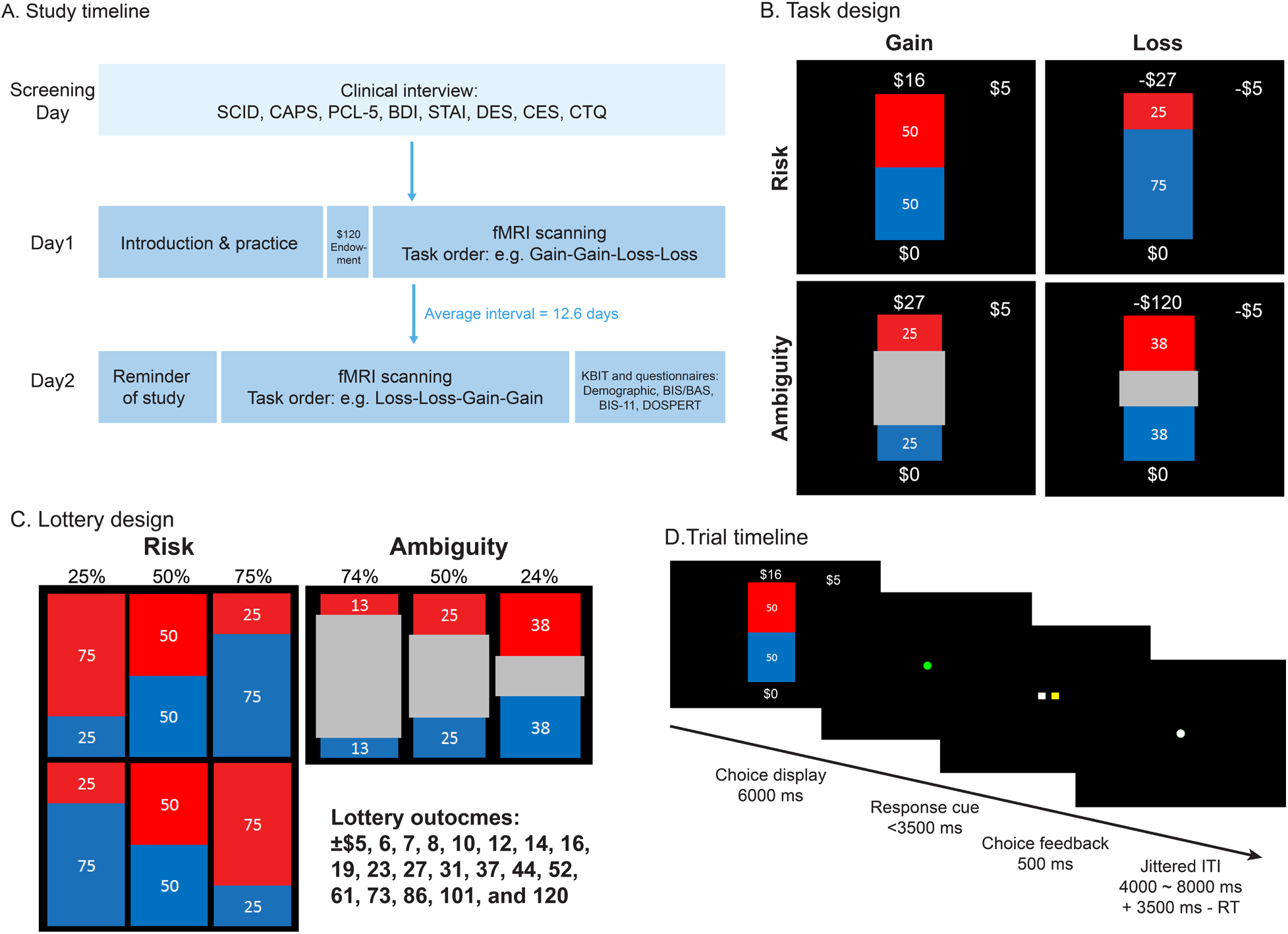
Study design. A: Timeline of the study. Participants went through a screening session and two scanning sessions on three different days. The screening session determined participants’ eligibility based on PTSD diagnosis, combat exposure, and exclusion of other neurological disorders. Eligible participants were scanned on two separate days on a decision making task. Measure labels: SCID: Structured Clinical Interview for DSM-4, CAPS: Clinician Administered PTSD Scale, PCL5:, PTSD Checklist for DSM-5, BDI: Beck Depression Inventory, STAI-1: State Anxiety, STAI-2: Trait Anxiety, DES: Dissociative Experiences Scale, CES: Combat Exposure Scale, CTQ: Childhood Trauma Questionnaire, KBIT: Kaufman Brief Intelligence Test, BIS/BAS: Behavioral Avoidance/Inhibition Scale, BIS-11: Barratt Impulsiveness Scale, DOSPERT: Domain-Specific Risk-Taking Scale. B: Task design: participants chose between a lottery and a sure outcome under four conditions: risky gains, ambiguous gains, risky losses, and ambiguous losses. Lotteries are shown as examples. Outcome probability of the risky lottery was represented by the area of the red or blue rectangle and was fully known to the participant. Outcome probability of the ambiguous lottery was covered by a grey rectangle in the middle, thus was partially known to the participant. C: Levels of risk (outcome probability, 0.25, 0.5, and 0.75), ambiguity (grey area, 0.74, 0.5, and 0.24), and monetary outcomes (20 monetary gains and 20 monetary losses) of the lottery. D: On each trial, participants had 6 seconds to view the options, and made a choice following a green response cue. They had a time limit of 3.5 seconds to register the choice, after which they would immediately see a confirmation with the yellow square representing the side they chose. The lottery was not played out during the scan to avoid learning. The inter-trial-interval (ITI) was jittered among 4, 6, and 8 seconds, and the remaining time during the response window (3.5 seconds – response time) would be added to the ITI.

### Clinical symptom variation

Participants varied in their PTSD symptom severity (Fig 2A) assessed by the Clinician-Administered PTSD Scale (CAPS) for DSM-4 (Diagnostic and Statistical Manual of Mental Disorders, 4^th^ Edition) (23). Veterans with PTSD had higher total CAPS score compared to controls (PTSD, N = 23: *Mean* = 72.13, *SD* = 15.04; control, N = 34: *Mean* = 6.21, *SD* = 9.68; *t*(34) = 18.58, *p* < 0.001). PTSD symptoms as captured by the 5-factor model of CAPS (1) were highly correlated with symptoms of depression, anxiety and dissociative experiences (Fig 2B, see Table S1 for descriptive statistics of all measures). In order to account for the influence of clinical symptoms on the behavior and neural activity during the task, we conducted principal component analysis (PCA) on these clinical symptoms. Since the severity of psychopathology may be affected by the degree of stress exposure, we also included measures of combat exposure (CES) and childhood trauma (CTQ) in the PCA. The first three components accounted for ~80% of the variance in those data (Fig S1A). The first component was affected by all clinical symptoms (PTSD, depression, anxiety, and dissociative experiences) and might reflect a general affective factor. This component was highly consistent with PTSD symptom severity (correlation with CAPS Spearman’s *ρ* = 0.94, *n* = 55, *p* < 0.001), and could be used to clearly classify PTSD diagnosis (Fig S1C). The second component was mostly affected by re-experiencing, avoidance and anxious arousal clusters of CAPS, as well as the degree of combat exposure, potentially representing a fear learning-updating deficit or general hyperarousal. The third component was affected by combat exposure and childhood trauma. Components 2 and 3 were not strongly correlated with PTSD symptom severity (n = 55, Component 2: correlation with CAPS Spearman’s *ρ* = 0.11, *p* = 0.43; Component 3: correlation with CAPS Spearman’s *ρ* = 0.029, *p* = 0.84).

**Figure 2.**
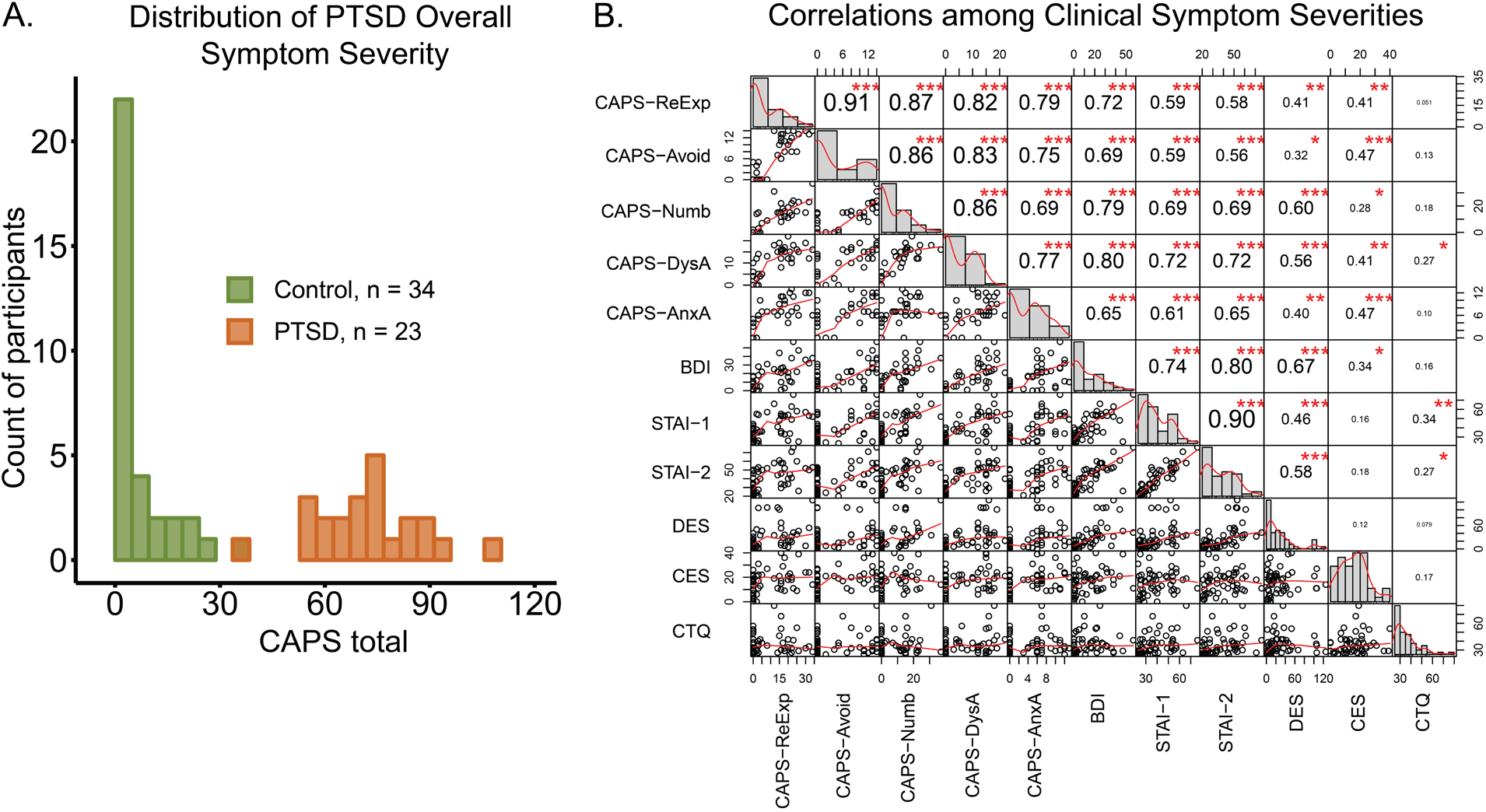
Participants’ symptom severity. A: Distribution of CAPS total score, colored by group (combat veterans with or without PTSD diagnoses). One PTSD participant included in the analysis did not have complete CAPS data. B: PTSD, depression and anxiety symptom severities were highly correlated. Numbers in the upper right panels indicate pair-wise Pearson correlation coefficients. Significance levels: ***, p < 0.001; **, p < 0.01; *, p < 0.05. Lower left panels show pairwise scatter plots and smoothed curves using locally weighted polynomial regression. Panels in the diagonal show distributions and density curves for each measure. Labels of measures: CAPS-ReExp: re-experiencing, CAPS-Avoid: avoidance, CAPS-Numb: numbing, CAPS-DysA: dysphoric arousal, CAPS-AnxA: anxious arousal, BDI: Beck Depression Inventory, STAI-1: State Anxiety, STAI-2: Trait Anxiety, DES: Dissociative Experiences Scale, CES: Combat Exposure Scale, CTQ: Childhood Trauma Questionnaire.

### PTSD symptom severity is associated with increased ambiguity aversion in the loss domain, and increased risk aversion in the gain domain

For each participant, we estimated risk and ambiguity attitudes for gains and losses, using the combined data from both scanning sessions (see equations 1 and 2 in Methods; see Fig S2A for an example from one participant). We then investigated the associations between these attitudes and PTSD diagnosis status, as well as PTSD symptom severity. All attitudes were transformed such that negative numbers indicate aversion to risk or ambiguity, and positive numbers indicate seeking. Based on the previous behavioral finding that PTSD symptom severity was associated with higher aversion to ambiguity in losses (6), we first investigated ambiguity attitudes. At the group level, participants were not significantly averse to ambiguity in the domain of losses (Fig 3A; PTSD: *Mean* = −0.25, *t*(23) = −1.81, *p* = 0.11; Control: *Mean* = 0.003, *t*(33) = 0.040, *p* = 0.97), and were significantly averse to ambiguity in the domain of gains (Fig 3A; PTSD: *Mean* = −0.35, *t*(23) = −3.45, *p* < 0.01; Control: *Mean* = −0.42, *t*(33) = −7.27, *p* < 0.001). However, a two-way ANOVA of ambiguity attitude with domain as the within-subject factor and group as the between-subject factor showed a significant interaction between domain and group (*F*(1,56) = 4.34, *p* < 0.05, *η^2^* = 0.0279). Post-hoc comparisons showed that veterans with PTSD were marginally more averse to ambiguity under losses (*p* = 0.081), but not under gains (*p* = 0.53). A dimensional analysis (Fig 3B) of this symptom–behavior relationship, regardless of PTSD diagnosis, revealed a negative correlation between ambiguity attitudes in the loss domain and CAPS total score (Spearman’s *ρ* with CAPS total score = −0.30, *p* < 0.05), indicating that higher symptom severity was related to higher aversion to ambiguity under losses. Since many control participants had a CAPS score of zero, we also repeated the analysis using PCL-5 scores instead of CAPS and overserved a similar effect (Fig S2B, Pearson’s *r* with PCL-5 = −0.31, *p* < 0.05).

**Figure 3.**
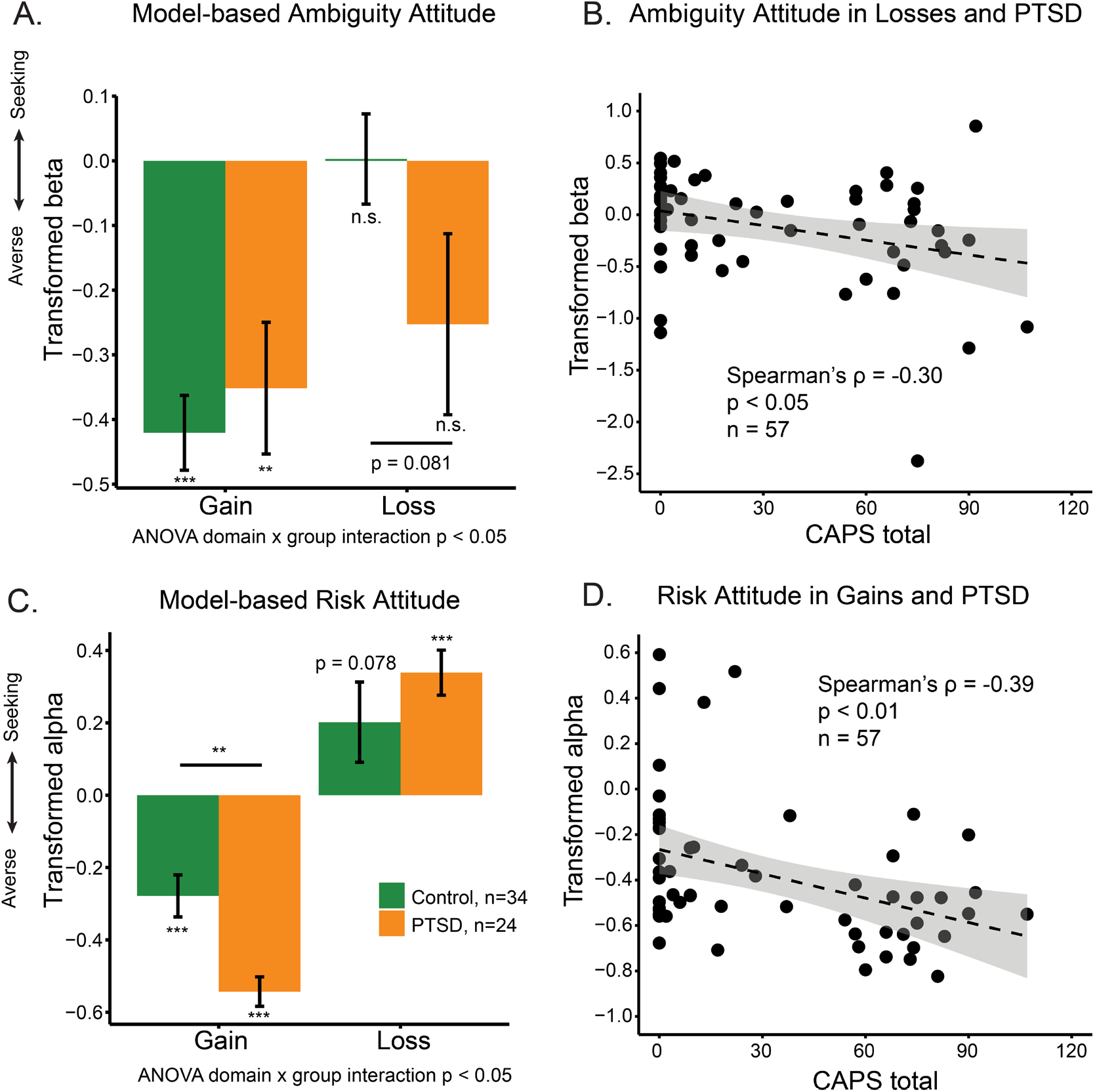
Uncertainty attitudes and PTSD symptom severity. A: Group comparison of ambiguity attitudes in gains and losses between veterans with PTSD and combat controls. B: PTSD symptom severity was negatively correlated with ambiguity attitude in losses. One participant was not included in the analysis due to missing CAPS. C: Group comparison of risk attitudes in gains and losses between veterans with PTSD and combat controls. D: PTSD symptom severity was negatively correlated with risk attitude in gains. One participant was not included in the analysis due to missing CAPS. In A and C, comparisons of each group’s attitudes with zero were FDR-corrected across all four comparisons in each uncertainty type. Post-hoc comparisons between groups in A and C are FDR-corrected. Significance level: *, p<0.05; **, p<0.01; ***, p<0.001.

Next, we examined risk attitudes. Both the PTSD and control groups exhibited risk aversion in the domain of gains (PTSD: *Mean* = −0.54, *t*(23) = −13.34, *p* < 0.001; Control: *Mean* = −0.28, *t*(33) = −4.80, *p* < 0.001). In the domain of losses, veterans with PTSD exhibited risk seeking (Fig 3C; PTSD: *Mean* = 0.34, *t*(23) = 5.43, *p* < 0.001), while combat controls exhibited marginal risk seeking (Control: *Mean* = 0.20, *t*(33) = 1.82, *p* = 0.078, FDR corrected for four comparisons). A two-way ANOVA of risk attitude with domain (gain or loss) as the within-subject factor and group as the between-subject factor revealed a significant interaction between domain and group (*F*(1,56) = 6.29, *p* < 0.05, *η^2^* = 0.0521). Post-hoc comparisons showed that veterans with PTSD were more averse to risk under gains (p < 0.01), but not under losses (*p* = 0.34), compared with combat controls. Examining this relationship further with a dimensional approach (Fig 3D and Fig S2C), we observed a similar effect: PTSD symptom severity was negatively correlated with risk attitudes in the gain domain (Spearman’s *ρ* with CAPS total = −0.39, *p* < 0.01; Pearson’s *r* with PCL5 = −0.36, *p* < 0.01).

Veterans with PTSD and combat controls did not differ in the choice noise parameter γ (a two-way ANOVA of γ with domain (gain or loss) as the within-subject factor and group as the between-subject factor: no main effect of group, *F*(1,56) = 1.28, *p* = 0.262, *η^2^* = 0.0120; no domain by group interaction, *F*(1,56) = 1.63, *p* = 0.207, *η^2^* = 0.0136). However, model-fitting quality was in general better in the control group than in the PTSD group (a two-way ANOVA of BIC with domain (gain or loss) as the within-subject factor and group as the between-subject factor: a main effect of group, *F*(1,56) = 4.75, *p* < 0.05, *η^2^* = 0.0587; no domain by group interaction, *F*(1,56) = 1.68, *p* = 0.200, *η^2^* = 0.00788).

To control for differences in age, income, education and intelligence, we used a linear regression model to explain uncertainty attitudes as a function of PTSD symptoms (CAPS total), while accounting for these demographic factors. Because model-fitting quality was affected by PTSD symptoms, we also included the BIC of the behavioral model as a predictor in the regression (see Supplementary Methods). For risk attitude in the gain domain, the effect of CAPS total score remained significant (multi-factor ANOVA by Generalized Linear Model: *F*(1, 41) = 12.5, *p* < 0.01). BIC was the only other significant factor (*F*(1, 41) = 17.7, *p* < 0.001). Similarly for ambiguity attitude in losses, CAPS total score (multi-factor ANOVA, *F*(1, 41) = 6.05, *p* < 0.05) and BIC (*F*(1, 41) = 4.86, *p* < 0.05) were the only significant factors.

Because seven of the combat-control veterans in this study sample also participated in the previous behavioral study, we also repeated the analysis excluding these returning participants, to yield a completely independent dataset. The negative relationships between PTSD symptom severity and ambiguity attitude in losses (Spearman’s *ρ* with CAPS total= −0.31, *p* < 0.05, *n* = 50), and between PTSD symptom severity and risk attitude in gains (Spearman’s *ρ* with CAPS total = −0.42, *p* < 0.01, *n* = 50) still held in this independent sample (Fig S3B, D).

We also assessed participants’ risk-taking attitudes through the Domain-Specific Risk-Taking (DOSPERT) Scale self-report questionnaire, but none of the domains (Ethical, Financial, Health/Safety, Recreational, and Social) was correlated with PTSD symptoms severity measured by CAPS total. Among the other self-report measures, CAPS total was correlated with total score of Behavioral Inhibition Scale (BIS, Spearman’s *ρ* = 0.37, *p* < 0.01, *n* = 57), and with total score of Barratt Impulsiveness Scale (BIS11, Spearman’s *ρ* = 0.47, p < 0.001, *n* = 57).

### PTSD symptom severity is associated with diminished neural response to decision making under uncertainty

To investigate the neural mechanisms of the stronger aversion to uncertainty observed in veterans with PTSD, we first examined the general neural activity during decision-making. Because the key process of our task was evaluating the subjective values of the uncertain options, we looked at the neural activity during the 6-second period of options presentation on each trial (see descriptive statistics of participants included in the neural analyses in Table S2). In a whole-brain analysis, we explored the relationship between PTSD symptom severity and the general neural activity during this valuation process (compared to baseline). Activity in a vmPFC - a central component of the valuation network - was negatively correlated with CAPS total score (*p* < 0.001, cluster-based corrected, Fig 4A), during the second session of the task. This negative relationship was not specific to a particular condition – rather, it was consistent across all four decision contexts (Fig 4B; Pearson’s *r*(risky gains) = −0.50, *r*(ambiguous gains) = −0.51, *r*(risky losses) = −0.51, *r*(ambiguous losses) = −0.40). Veterans with higher overall PTSD symptom severity showed more vmPFC deactivation during valuation of uncertain options. This finding is consistent with our hypothesis regarding the valuation system’s involvement in PTSD; next, we directly examined the neural correlates of valuation in the task.

**Figure 4.**
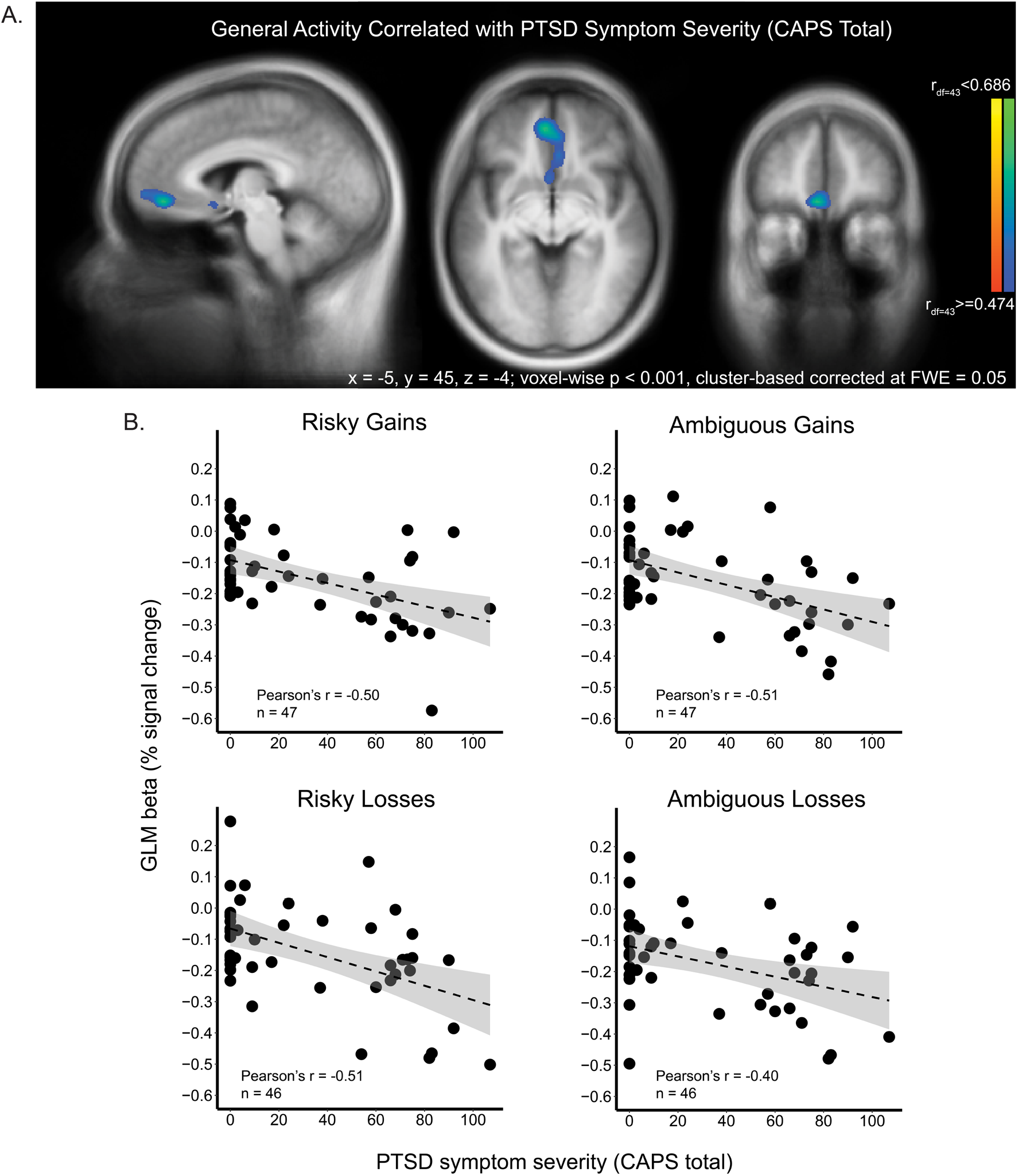
Reduced vmPFC activity during valuation is related to PTSD symptom severity. A: A whole-brain analysis revealed that activity in vmPFC during valuation was negatively correlated with CAPS total score, regardless of decision condition. B: Visualization of this negative correlation between general activity in the vmPFC and CAPS total score in each decision condition.

### PTSD symptom severity is associated with altered neural encoding of subjective value of uncertain options

For each participant, we calculated the subjective value of the lottery presented on each trial, based on the behavioral model (see equation 1 in Methods), using the participant-specific risk and ambiguity attitudes under gains and losses. We then included the subjective values (positive for gain lotteries, negative for loss lotteries) in the GLM, separately for each of the four decision conditions. We focused our analysis on the two decision conditions in which symptoms influenced behavior: ambiguous losses and risky gains. To examine group differences between veterans with PTSD and combat controls, we directly contrasted their neural representation of subjective value in a whole-brain analysis. Veterans with PTSD showed more negative subjective-value signals for ambiguous losses in left inferior frontal regions (IFG) and bilateral occipital regions, compared to controls (Fig 5A; for statistics of all regions, see Table S3). We then used a leave-one-subject-out (LOSO) procedure to define regions of interest around the inferior frontal gyrus (IFG) and sample activation in an unbiased manner (see Methods). The subjective-value signal of ambiguous losses in IFG was negatively correlated with PTSD symptom severity (Fig 5B; Spearman’s *ρ* = −0.35, *p* < 0.05, *n* = 48), such that higher symptom severity was associated with more negative subjective-value signal. Veterans with PTSD showed more positive subjective-value signals for risky gains in right orbitofrontal cortex (OFC) in a whole-brain analysis (Fig 5C), and PTSD symptoms severity was positively correlated with subjective-value signal of risky gains in this OFC region (Fig 5D; Spearman’s *ρ* = 0.52, *p* < 0.001, *n* = 48). For completion, we also looked at the other two conditions. Veterans with PTSD showed more positive encoding of subjective value of ambiguous gains in the thalamus and right cerebellum (Fig S4), and there was no group difference in the subjective-value encoding of risky losses.

**Figure 5.**
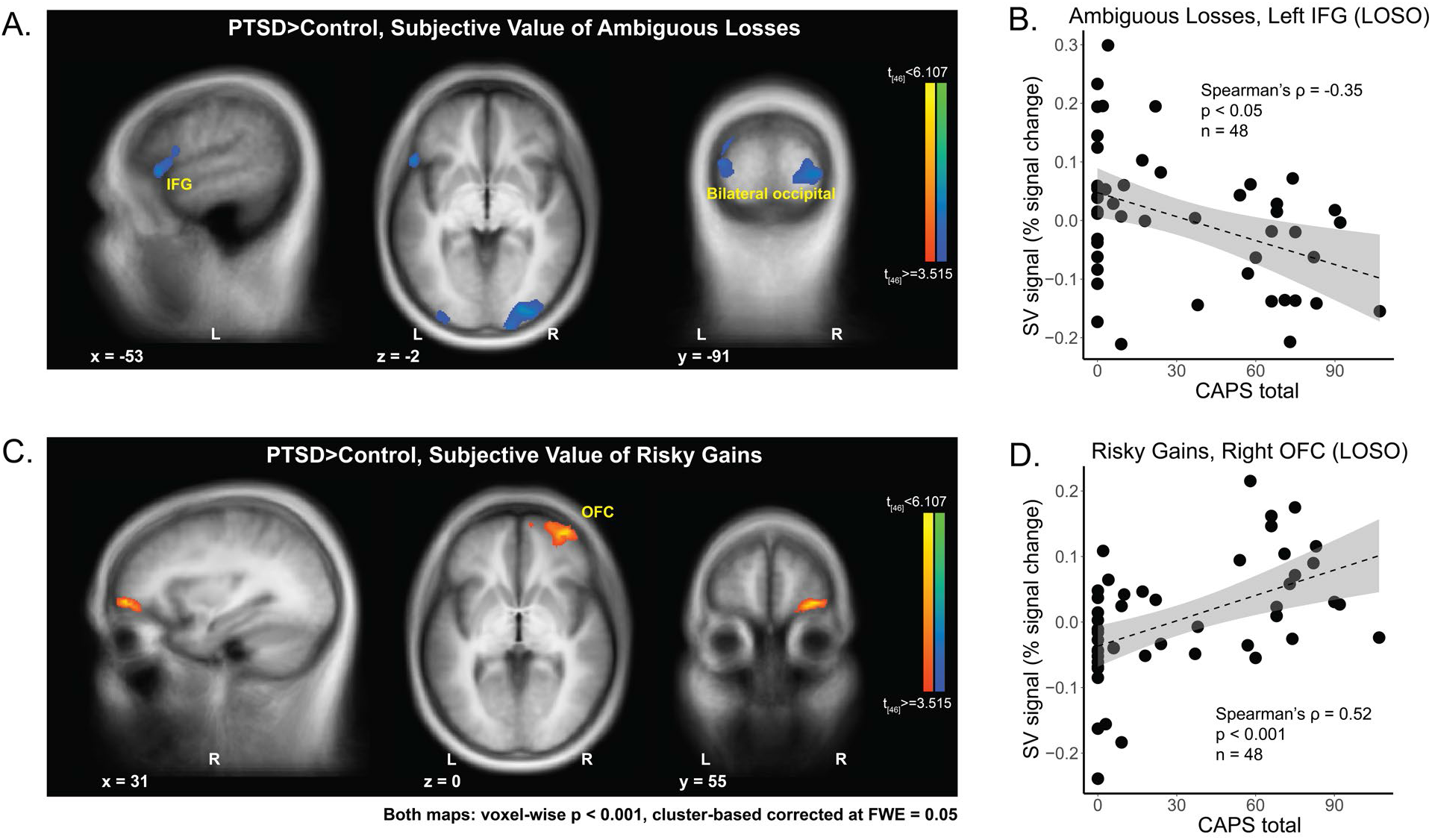
Neural representation of subjective value directly contrasting PTSD and control. Whole-brain comparisons of neural subjective-value signals between veterans with PTSD and combat controls, under A: ambiguous losses, and C: risky gains. All maps were corrected using cluster-based method controlling family-wise error at 0.05, when thresholded at p < 0.001 at the voxel level. B: Neural subjective-value representation of ambiguous losses in the left IFG was negatively correlated with PTSD symptom severity. D: Neural subjective-value representation of risky gains in the right OFC was positively correlated with PTSD symptom severity. ROIs in B and D were defined by a leave-one-subject-out (LOSO) approach.

To further probe group and individual differences in value encoding, we examined the subjective-value signals of each group in the classical value areas – the vmPFC and the ventral striatum – as defined in a meta-analysis by Bartra and colleagues (14). We again focused on the conditions of ambiguous losses and risky gains (Fig 6). In vmPFC, the subjective-value signal of risky gain lotteries was positively correlated with PTSD symptom severity (Fig 6A, Spearman’s *ρ* with CAPS = 0.31, *p* < 0.05). In ventral striatum, subjective-value signal of ambiguous loss lotteries was negatively correlated with PTSD symptom severity (Fig 6B, Spearman’s *ρ* with CAPS = −0.35, *p* < 0.05). PTSD symptom severity was not significantly associated with the subjective-value signal of ambiguous losses in vmPFC (Fig 6A, Spearman’s *ρ* with CAPS = −0.18, *p* = 0.22), or with the subjective-value signal of risky gains in ventral striatum (Fig 6B, Spearman’s *ρ* with CAPS = 0.22, *p* = 0.14; see Fig S5 A and B for correlations with PCL5). These relationships could also be revealed in the group comparison (Fig 6 C and D). The subjective-value signal of ambiguous losses was more negatively encoded in ventral striatum in veterans with PTSD compared with combat controls (Fig 6D, *t* = −2.77, *p* < 0.01). Conversely, the subjective-value signal of risky gains was marginally more positively encoded in vmPFC in veterans with PTSD than in combat controls (Fig 6C, *t* = 1.97, *p* = 0.054).

**Figure 6.**
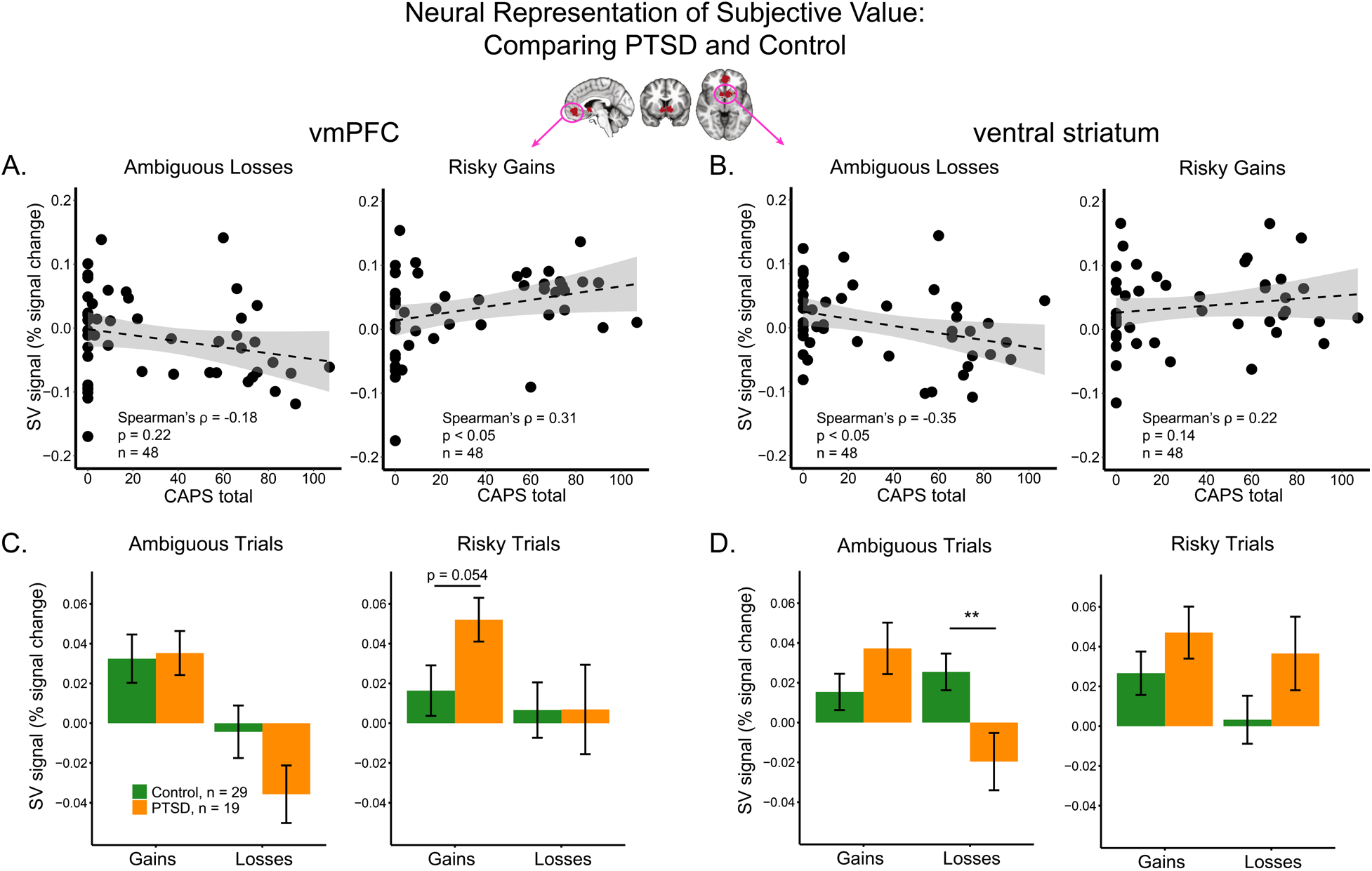
Neural subjective-value signals in external ROIs of vmPFC and ventral striatum were related to PTSD symptom severity. A: In vmPFC, correlations between subjective-value signals of ambiguous losses and risky gains and PTSD symptom severity (CAPS total). B: In ventral striatum, correlations between subjective-value signals of ambiguous losses and risky gains and PTSD symptom severity (CAPS total). C: In vmPFC, group comparison of neural subjective-value signals between veterans with PTSD and combat controls. D: In ventral striatum, group comparison of neural subjective-value signals between veterans with PTSD and combat controls. In panels C and D, comparisons were post-hoc FDR-corrected after ANOVA within each figure. Significance level: *, p<0.05; **, p<0.01; ***, p<0.001. ROIs of vmPFC and ventral striatum were taken from Bartra and colleagues’ meta-analysis study (14).

The relationships between subjective-value signals and PTSD symptom severity held after controlling for age, income, education and intelligence (see Supplementary Methods). The subjective-value signal of ambiguous losses in ventral striatum was affected by CAPS (multi-factor ANOVA by Generalized Linear Model, *F*(1, 33) = 6.01, *p* < 0.05), and not by the four demographic factors. The subjective-value signal of risky gain lotteries in vmPFC was marginally affected by CAPS (multi-factor ANOVA by Generalized Linear Model: *F*(1, 33) = 3.53, *p* = 0.069), and not by the four demographic factors.

### A shift from value-encoding to saliency-encoding of ambiguous losses in PTSD

Our results so far point to differences in the mechanisms of subjective-value encoding between veterans with PTSD and combat controls. This difference is most notable for ambiguous losses: in combat controls, ambiguous losses were encoded in a positive manner (decreased activity for increased losses) consistent with a monotonic representation of value. Conversely, in the brains of veterans with PTSD, losses were encoded negatively (increased activity for increased losses), consistent with a U-shaped saliency-encoding mechanism. This difference in representation was particularly striking in the ventral striatum (Fig 6D). To directly confirm this group difference, however, we need to examine gains and losses on the same scale. To this end, we constructed a GLM with one predictor for the value of ambiguous gains and losses, and another predictor for the saliency of the same gains and losses. Subjective values of the lotteries were used for the value predictor, and saliency was computed as the absolute value of these subjective values (Fig 7A; see Methods for fMRI GLM first-level analysis). While the ventral striatum in controls significantly encoded value (one-sample t test GLM beta compared with 0, *t*(28) = 3.4, *p* < 0.01), but not saliency (*t*(28) = −0.62, *p* = 0.54), the opposite pattern was observed in veterans with PTSD: activity in the same brain area in the PTSD group encoded saliency (*t*(18) = 2.7, *p* < 0.05), but not value ((*t*(18) = 0.99, *p* = 0.45; all *p* values were FDR corrected for four comparisons). Furthermore, the saliency-encoding patterns were significantly different between veterans with PTSD and combat controls (two-sample t test: *t*(39.3) = −2.5, *p* < 0.05).

**Figure 7.**
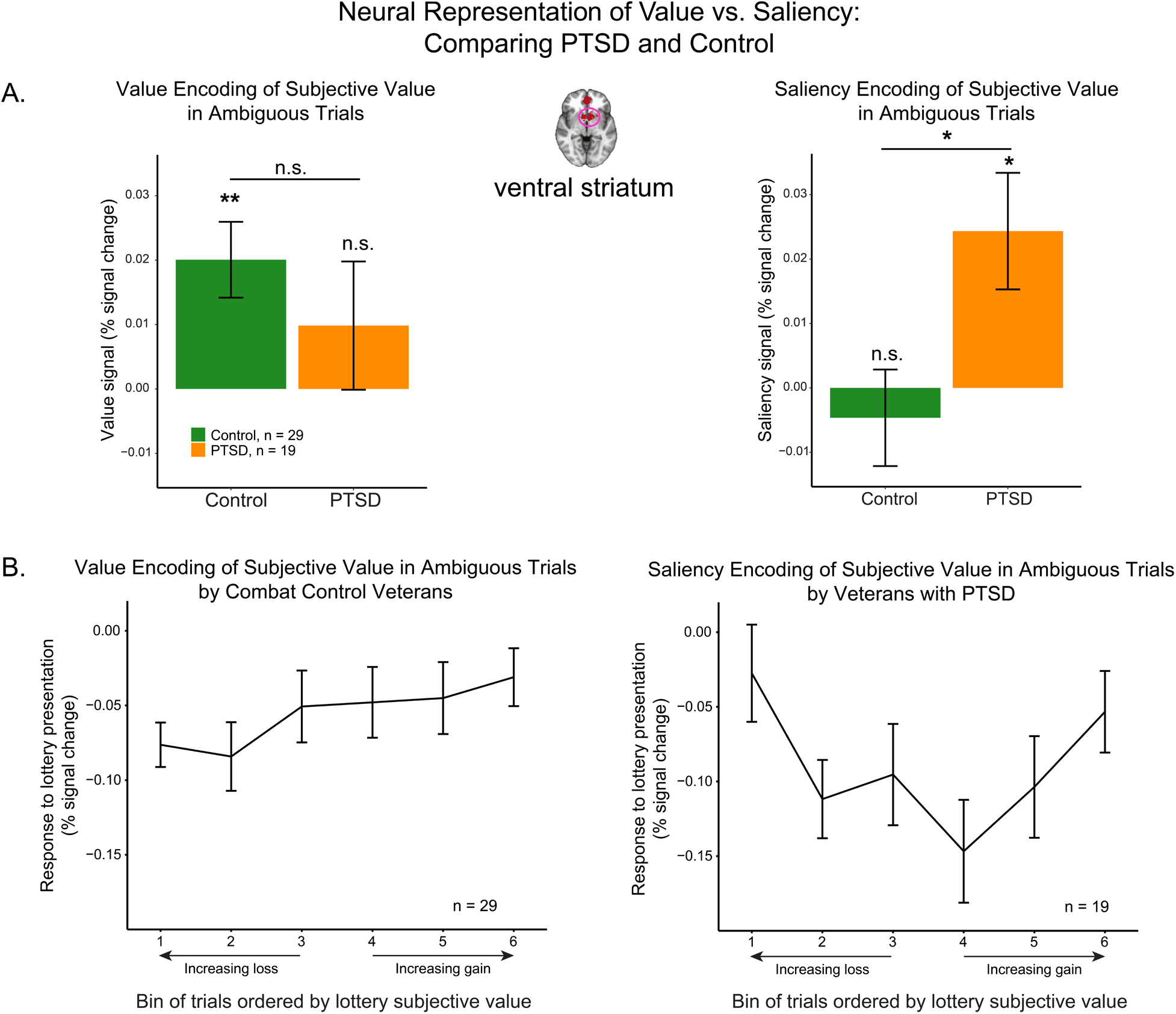
Value and saliency encoding in the ventral striatum of PTSD and controls. A: In ventral striatum, value-encoding of subjective values was observed in combat controls but not in veterans with PTSD; saliency-encoding of subjective values was observed in veterans with PTSD but not in combat controls. Comparisons with zero for both PTSD and Control group were FDR-corrected across four comparisons in the two figures. Significance level: *, p<0.05; **, p<0.01; ***, p<0.001. B: Direct visualization of neural response to trials of ambiguous lotteries with different levels of subjective values in ventral striatum. Bins were ordered monotonically based on participant-specific subjective values of the lotteries across losses and gains. Bins 1-3 were loss lotteries, and bins 4-6 were gain lotteries. Consistent with panel A, combat control veterans encoded subjective value in a monotonic value-pattern, and veterans with PTSD encoded subjective value in a U-shaped saliency-pattern. ROI of ventral striatum was taken from Bartra and colleagues’ meta-analysis study (14). All error bars indicate standard errors.

Figure 7B presents a direct visualization of the shape of value- and saliency-encoding in the ventral striatum in the two groups of veterans. For each participant, and within each uncertainty domain (risk or ambiguity), we grouped the trials into 6 bins across losses and gains, based on the subjective values of the lotteries. The 1^st^ bin corresponded to the loss lotteries with the most negative subjective values, and the 6^th^ bin corresponded to the gain lotteries with the most positive subjective values (see Methods for details). As expected, combat control veterans showed a monotonic representation of subjective value, whereas veterans with PTSD showed a U-shaped representation (Fig 7B).

### Neural activity explains symptoms variation better than behavioral uncertainty attitudes

Having revealed both behavioral and neural alteration related to PTSD, we explored whether PTSD symptom variation could be better explained by neural activation patterns, behavioral uncertainty attitudes, or a combination of both. We constructed three linear models to predict PTSD symptom severity indicated by CAPS total score, using (1) magnitudes of neural responses to the four decision conditions in vmPFC (ROI from Fig 4); (2) behavioral risk and ambiguity attitudes under gains and losses; and (3) both neural responses and behavioral attitudes. All models controlled for age and intelligence (KBIT) (see details in Supplementary Methods). The model including only neural measures best explained the variation of PTSD symptom severity (BIC(neural model) = 132.8, BIC(behavioral model) = 154.7, BIC(full model) = 143.8, Fig S6).

### Emotional numbing plays the key role in diminished vmPFC general neural activity

So far, we investigated the relationship between PTSD overall symptom severity and valuation under uncertainty. We further looked at whether a specific symptom cluster was the main source of influence, considering the multi-dimensional nature of PTSD. From the vmPFC ROI in which neural activity was negatively correlated with CAPS total score in the whole-brain analysis (Fig 4A), we sampled general neural activity during the valuation phase of the task (GLM beta) from each participant. We then constructed a linear regression model to explain this region’s activity with all five clusters of CAPS, including re-experiencing, avoidance, emotional numbing, dysphoric arousal, and anxious arousal, accounting for age and intelligence. Only the emotional numbing cluster significantly contributed to the negative correlation between vmPFC activity and PTSD symptom severity (standardized regression coefficient, *Beta* = −0.72, *t* = −2.32, *p* < 0.05, Fig S7A). Age and intelligence (KBIT) did not significantly influence vmPFC neural activity (standardized regression coefficient, Age: *Beta* = −0.14, *t* = −1.13, *p* = 0.26; intelligence: *Beta* = −0.044, *t* = 0.33, *p* = 0.75). Variable selection using exhaustive search also indicated that including only the emotional numbing cluster out of all PTSD symptom clusters best explained the relationship between vmPFC neural activity and PTSD symptom severities (Fig S7B, BIC = 112.8; see details in Methods for fMRI GLM second-level analysis).

### Influence of clinical symptoms beyond PTSD

Because our participants showed high levels of comorbidity with other clinical symptoms, especially depression and anxiety (Fig 2B), we also investigated how the behavioral and neural mechanisms of valuation were influenced by symptoms beyond PTSD. We examined the correlation between the first three principal components of all clinical measures (Fig S1) and the behavioral uncertainty attitudes. Principal component 1 (general affective symptom) was negatively correlated with risk attitude under gains (Pearson’s *r* = −0.35, *p* < 0.01) and ambiguity attitude under losses (Pearson’s *r* = −0.29, *p* < 0.05), consistent with the effect of the overall PTSD severity indicated by CAPS total score (Fig 3). We did not find any relationship between uncertainty attitudes and the second (fear learning-updating) or the third (trauma severity) principal components.

We also examined potential relationships between subjective-value signals and the three principal components. In vmPFC, the first component (general affective symptom) was positively correlated with encoding of subjective value of risky gains (Pearson’s *r* = 0.30, *n* = 47, *p* < 0.05), consistent with the effect of PTSD symptom severity (CAPS total score). The third component (trauma severity) was negatively correlated with encoding of subjective value of ambiguous losses (Pearson’s *r* = −0.29, *n* = 47, *p* < 0.05), in the same direction as the correlation between subjective value of ambiguous losses and PTSD symptom severity. In ventral striatum, no correlation survived our statistical thresholds (see all correlations in Fig S5C).

## Discussion

In this study, we explored the neural basis of valuing uncertain monetary rewards and punishments, in veterans exposed to combat trauma with a wide range of PTSD symptoms. Behaviorally, symptom severity was associated with increased aversion to ambiguous losses, and increased aversion to risky gains. These two conditions were also the ones in which PTSD symptom severity influenced the neural representations of subjective value (Fig 7). Two main effects were observed: first, in both whole-brain and ROI analyses, veterans with PTSD showed more negative neural representation of ambiguous losses, and more positive neural representation of risky gains, than combat control veterans. Second, there was a qualitative group difference in the neural representation of ambiguous lotteries in the ventral striatum. In veterans with PTSD, this region encoded the saliency of the lotteries (with increased activity for both potential large gains and potential large losses), whereas in combat control veterans it encoded the lottery value. Moreover, a direct examination of the neural response to varying subjective values (Fig. 7B) suggests that the value pattern in controls was weak compared to the saliency pattern in PTSD. An intriguing possibility is that the strong neural tracking of saliency is a marker for vulnerability to PTSD, reflecting increased sensitivity to highly salient stimuli. The value signal, on the other hand, may be a marker of resiliency to PTSD. Future research, and in particular longitudinal studies that compare individuals exposed to trauma to those who never experienced trauma, are needed to explore this possibility.

### Using behavioral economics to identify markers of psychopathology

Our results add to a growing body of research, demonstrating the utility of behavioral economics in studying psychopathology (24–28). Replicating the previous behavioral study (6), we found an association between higher PTSD symptom severity and greater ambiguity aversion under losses, in an independent combat veteran sample. It should be noted, however, that this effect is weak, and was not significant in the group comparison. A larger sample is needed to further confirm the robust effect of the relationship. We also identified greater aversion to risk under gains in veterans with PTSD, likely due to a task design with increased range and variance of monetary outcomes that provided higher sensitivity for capturing true uncertainty attitudes. Our neural measure allowed us to also quantify individual and group differences in neural sensitivity to rewards and punishments. Previous studies have shown alterations in the neural processing of aversive outcomes in individuals with PTSD in various brain areas, including several medial and lateral prefrontal regions. Many of these studies, however, used fear and trauma-related stimuli (29). Here we show that activation in the same brain areas is affected by PTSD symptoms even in an economic decision task, completely unrelated to the trauma. This raises the possibility of developing diagnostic methods in the domain of decision making under uncertainty, which do not require patients to recall the traumatic experience. Several previous studies have also reported altered reward processing in PTSD (30), including reduced expectation of uncertain monetary outcomes (31,32) and decreased differentiation between monetary gains and losses in the striatum (22). Our experimental approach allowed us to estimate individual uncertainty attitudes during active decision making under four unique contexts. We applied a well-established computational model to infer these behavioral individual differences from the observed choice behavior (rather than estimating them through self-reports) and used the individual differences in the analysis of the neural data. Interestingly, participants’ self-reported risk-taking on the DOSPERT questionnaire was not strongly correlated with their PTSD symptom severity, suggesting that our method for estimating uncertainty attitudes through a behavioral task may be more sensitive for capturing subtle differences associated with clinical symptoms. An intriguing question remains to be answered is what contributes to the context-specific differences associated with PTSD symptom severity. We found higher behavioral aversion only to ambiguous losses and risky gains in PTSD, and the most striking neural difference of subjective-value encoding was revealed in the context of ambiguous losses. Compared with risky outcomes, of which both outcome magnitude and probability are known, ambiguous outcomes lack the exact information of outcome probability. This additional level of uncertainty may be more relevant to the nature of battlefield, making negative ambiguous outcomes more relatable to combat exposure. Our task design separating decision contexts enabled us to pinpoint specific cognitive processes affected by PTSD in combat veterans, but replication in larger samples is needed to confirm the results.

### Neural processing of rewards and punishments is associated with PTSD symptoms

By including both monetary gains and losses in the task design, we identified a shift from value-encoding to saliency-encoding in the brains of individuals who developed PTSD following trauma exposure (Fig 7). This shift could potentially imply an attention or arousal signal, that leads to avoidance of aversive outcomes like uncertain monetary gains or losses. Several previous studies examined the neural processing of value and saliency and revealed both distinct and overlapping regions for each type of encoding. Value signals were found in ventral striatum, parietal cortex, OFC, rostral ACC, and saliency signals were found in ventral striatum, rostral ACC, dorsal ACC, anterior insula by both univariate and multivariate analyses (12,13,15–18,33,34). To our knowledge, our results are the first to recognize the influence of psychiatric symptoms in humans on the value/saliency-encoding pattern. Interestingly, recent research in mice shows a similar flip in representation, where acute stress transforms reward responses in the lateral habenula into punishment responses (35). Neurons in the nucleus accumbens of rats can also flexibly shift their preferences between rewards and punishments, based on the emotional environment (36), suggesting that what we observe here may reflect a stress coping mechanism.

PTSD is highly comorbid with symptoms of depression and anxiety. Through PCA, we were able to disentangle three main symptom components, and showed that the component of general affective symptoms was likely the main source of influence (Fig S5). In addition, we also found that the component of trauma symptoms was related to the neural representation of subjective value of ambiguous losses in vmPFC, raising the possibility that trauma exposure additionally influences sensitivity to aversive monetary outcomes, independent from general affective symptoms. Trauma symptoms were assessed through combat exposure and childhood trauma in our sample of veterans. Future research could investigate more systematically how other types of trauma exposure could additionally influence the neural processing of valuation of uncertain outcomes.

One concern in our investigation of neural representation of value is that the range of subjective values is lower in the group of veterans of PTSD because of their higher aversion to uncertainty, which could influence the sensitivity of the neural response to value differences. It should be noted, however, that our main conclusion is based on a difference in the direction of correlation (negative vs. positive), rather than a difference in the magnitude of slope of the correlation (Figs 6 and 7). This represents a substantial difference in the shape of subjective-value encoding and would not be affected by group difference in the range of subjective values.

### Neural markers of vulnerability and resilience to PTSD

Previous studies of PTSD often focused on the neural processing of fear and trauma, and identified both functional and structural abnormalities in amygdala, hippocampus, and vmPFC (5,29,37–39). Other studies have looked into more general cognitive processes and found blunted neural activation to monetary rewards (21,22). In our study, using a more nuanced computational approach, PTSD symptoms were associated with increased neural sensitivity to rewards and opposite direction of sensitivity to punishments. While the sensitivity to rewards may seem at odds with the previous studies, it should be noted that in those studies reward signals were defined as the difference in activation to gains and losses. A weaker contrast in individuals with PTSD could stem from a weaker reward signal, but also from a stronger punishment signal, consistent with a U-shaped saliency representation, as we report here, in which both highly salient positive and highly salient negative outcomes elicit similar magnitude of neural activation. With this being said, in the reward domain, we do find evidence of surprisingly stronger value sensitivity in PTSD (Fig 5 and 6). Note that reduced activation to rewards in individuals with PTSD was previously observed in comparison to controls who *were not exposed to trauma* (21). In our study, veterans with PTSD exhibited neural patterns for potential rewards which were similar to what has been observed in the general population (9,14). Combat controls, who were exposed to trauma, but did not develop PTSD, were the ones who differed from the general population, suggesting that they have exhibited a resiliency marker. Other important methodological details may contribute to the difference in results between the studies, including a focus on outcome delivery, rather than decision value, female participants, and mixed types of trauma. These differences suggest interesting directions for future research. Of particular interest is the interaction between uncertainty and combat trauma, as uncertainty is a central component of the battlefield experience. Comparing behavior and neural mechanisms in individuals who experienced combat trauma and those who did not, will help to shed light on the potential vulnerability and resiliency markers proposed here.

In line with the NIMH Research Domain Criteria (RDoC), we did not exclude veterans with history of substance abuse, to allow for a diverse representative sample of trauma exposed symptomatology. We controlled for substance abuse by conducting urine test and breathalyzer for anyone with substance abuse history or if we suspected any intoxication, and excluded those with positive results. The severity for substance abuse history in our sample was low and did not vary too much as measured by the Addiction Severity Index (ASI-alcohol: median = 0.089, range = [0, 1.47]; ASI-drug: median = 0, range = [0, 0.092]). Future research could better control for substance abuse history and medication, and potentially look into the pharmacological effect involving the dopamine and serotonin systems, which are crucial for value-based decision making (40,41).

Our study could not establish causal relationship between decision making under uncertainty and the development of PTSD symptoms. Heightened aversion to uncertainty could possibly predispose individuals to developing PTSD symptoms, and on the other hand, acquiring PTSD symptoms could result in altered uncertainty attitudes. There is some evidence, however, that risk attitude is correlated with relatively stable biomarkers including structural volume of right posterior parietal cortex (42), structural and functional connectivity of the amygdala (43) and genetic variations (44). These pieces of evidence might indicate that risk attitude is a personal trait, raising the possibility of its predisposing effect on the development of PTSD symptoms. Less evidence exists for biomarkers of ambiguity attitude, although there is some evidence for a genetic association among females (45). Further longitudinal studies comparing veterans pre- and post-military service may disentangle the role of pre-existing uncertainty attitudes on the development of PTSD from the subsequent impact of PTSD symptomatology on uncertainty attitudes.

Variations in decision making under uncertainty, and especially under ambiguity, have also been reported in other psychiatric disorders, including higher ambiguity aversion and choice inconsistency in individuals with Obsessive Compulsive Disorder (25), and decreased ambiguity aversion in individuals with antisocial personality disorder (26). Interestingly, a recent longitudinal study demonstrated transient increases in tolerance to ambiguity before relapses in opioid users undergoing treatment (27). Overall, these efforts to study psychiatric disorders using behavioral economics approaches could collectively lead to both early identification of behavioral and biological risk factors for symptom development, and more effective treatment.

## Methods

### Participants

68 male veterans (ages: 23.6-74.6; mean ± standard deviation: 39.4 ± 11.5), who had been deployed and exposed to combat, were recruited through flyers and were screened by clinicians at West Haven Veterans Affairs hospital. Due to the small proportion of female combat veterans (15% of female in Army 2019, statistics from Department of Defense), we only included male participants. PTSD sympotms and diagnoses were determined by the Structured Clinical Interview for DSM-4 (SCID) (46) and the Clinician Administered PTSD Scale (CAPS) (23). Participants either had current diagnoses of PTSD at the time of the study or were never diagnosed with PTSD (controls). We also collected the following measurements: PTSD Checklist for DSM-5 (PCL-5) (47), Beck’s Depression Inventory (BDI) (48), State-Trait Anxiety Inventory (STAI) (49), Dissociative Experiences Scale (DES) (50), Combat Exposure Scale (CES) (51), and Childhood Trauma Questionnaire (CTQ) (52). Participants with psychosis, bipolar disorder, traumatic brain injury, neurologic disorder, learning disability, and ADHD were excluded after screening. Participants also completed other questionnaires including demographic information, Behavioral Avoidance/Inhibition (BIS/BAS) Scales (53), the Barratt Impulsiveness Scale (BIS-11) (54), and Doman-Specific Risk-Taking (DOSPERT) Scale (55). Kaufman Brief Intelligence Test (KBIT) (56) was administered after scanning as a measure of non-verbal intelligence.

Participants data was excluded based on behavioral quality check (see Supplementary Methods) and excessive movement in the scanner. Behavioral data of 58 participants (ages: 23.6-67.0; mean ± standard deviation: 37.3 ± 8.9) and neural results of 48 participants (ages: 23.6-67.0; mean ± standard deviation: 37.4 ± 9.2), were reported. The study was approved by the Yale University Human Investigating Committee and the Human Subjects Subcommittee of the VA Connecticut Healthcare System, and compliance with all relevant ethical regulations was ensured throughout the study. All participants gave informed consent and were compensated with $100 for their participation, plus a variable bonus ($0-$240) based on choices they made in the task (see Supplementary Methods).

### Experimental design

The study was composed of three separate visits on three different days (Fig 1A). On the first day, recruited participants went through clinical interviews for screening. Eligible participants continued to two fMRI sessions, on two separate days. In the scanner, participants performed a task of decision making under uncertainty, which is based on a previous neuroimaging study (57) and similar to the design of a previous behavioral study in combat veterans (6). They made a series of decisions between a sure monetary outcome and an uncertain monetary outcome with either known (risky) or unknown (ambiguous) outcome probability, in scenarios of both gaining and losing money (Fig 1B). On each trial, participants viewed the two options side-by-side for a fixed duration of 6 seconds, and then made a choice (Fig 1D). To prevent learning, the outcome of the chosen option was not presented during the scan. At the end of the experiment, one randomly selected trial was realized for bonus payment. The scans were conducted over two days in order to limit the scanning time in each visit; task designs were identical for scanning Day1 and Day2. Participants were introduced to the task at the beginning on the Day and were reminded of the study on Day2. Additional questionnaires and intelligence tests were administered at the end of Day2.

### MRI scans

MRI data were collected with two scanners (due to scanner upgrade) at the Yale Magnetic Resonance Research Center: Siemens 3T Trio (37 participants, 29 reported in imaging results) and 3T Prisma (31 participants, 19 reported in imaging results), using a 32-channel receiver array head coil. High resolution structural images were acquired by Magnetization-Prepared Rapid Gradient-Echo (MPRAGE) imaging (TR = 2.5 s, TE = 2.77 ms, TI = 1100 ms, flip angle = 7°, 176 sagittal slices, voxel size = 1 ×1 × 1 mm, 256 × 256 matrix in a 256 mm field-of-view, or FOV). Functional MRI scans were acquired while the participants were performing the choice task, using a multi-band Echo-planar Imaging (EPI) sequence (TR= 1000 ms, TE= 30ms, flip angle=60°, voxel size = 2 × 2× 2 mm, 60 2 mm-thick slices, in-plane resolution = 2 × 2 mm, FOV= 220mm).

### Model-based risk and ambiguity attitudes estimation

We fitted each participant’s choice data separately into a behavioral economics model that was used in previous studies (6,9). The model fitting was conducted separately for gain and loss trials. The model separates the decision process into two steps: valuation and choice. In the valuation step, the subjective value (SV) of each option is modelled by equation (1),

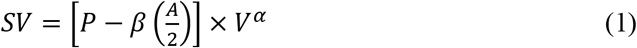

where P is the outcome probability (0.25, 0.50, or 0.75 for risky lotteries, 0.5 for ambiguous lotteries, and 1 for the certain option); A is the ambiguity level (0.24, 0.5, or 0.74 for ambiguous lotteries; 0 for risky lotteries and the certain amount); V is the non-zero outcome magnitude of the lottery or the amount of money of the certain option. For choices in the loss domain, amounts are entered with a positive sign. Risk attitude was modeled by discounting the objective outcome magnitude by a participant-specific parameter, α. In the gain domain, a participant is risk averse when α < 1, and is risk seeking when α > 1. Because we fitted the choice data in the loss domain using positive outcome magnitudes, the participant is risk averse when α > 1, and is risk seeking when α < 1. Ambiguity attitude was modeled by discounting the lottery probability linearly by the ambiguity level, weighted by a second participant-specific parameter, β. A participant is averse to ambiguity when β > 0, and is ambiguity seeking when β < 0 in the gain domain. In the loss domain, participant is averse to ambiguity when β < 0, and ambiguity seeking when β > 0.

The choice process is modeled by a standard soft-max function (equation 2),

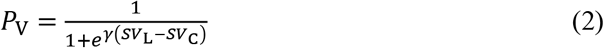

where PV is the probability of choosing the lottery option, SVC and SVL are the subjective values of the certain option and the lottery respectively, calculated by equation (1); γ is a participant-specific noise parameter. We fitted each participant’s choices combining data from two sessions and obtained four attitudes: risk attitudes for gains and losses, ambiguity attitudes for gains and losses. For consistency, we transformed all attitudes in the following way such that negative values indicate aversion and positive values indicate seeking: risky gains: α – 1, risky losses: 1 – α, ambiguous gains: - β, ambiguous losses: β. Since participants performed the task on two separate sessions, we also fitted each session’s choice data separately. These fitted parameters from separate sessions were used to calculate trial-wise subjective values of the lotteries for GLM neural analysis, because they could capture the subjective values more accurately for searching neural activity change induced by variations of subjective values.

### MRI data analysis

MRI data were preprocessed in BrainVoyager (Version 20.2.0.3065). Anatomical images were normalized to the standard brain template in Talairach space for each participant. Preprocessing of functional data included motion correction, slice scan time correction (cubic spline interpolation), temporal filtering (high-pass frequency-space filter with cut-off cycle of 3), spatial smoothing (Gaussian filter with 8mm full-width at half-maximum), co-registration with high-resolution standardized anatomical data, and normalization to Talairach space. Scan data with movement of over 2 mm in any direction were excluded from analysis.

First level GLM analysis was conducted in the Neuroelf toolbox (Version 1.0, https://neuroelf.net/) through MATLAB (Version R2018b). The pre-processed fMRI signal time course was first converted to percent signal change within each scanning block, and activity of each voxel was modeled by GLM predictors convolved with a standard double-gamma hemodynamic response function. In the first GLM, we looked at the general activity during decision making, by including four binary predictors for all four decision conditions: ambiguous gains, risky gains, ambiguous losses, and risky losses. Each binary predictor was modeled as a box-car function, with the duration of choice display (6TR). We modeled choice response of all trials by another binary predictor with the duration of 1TR at the time of button press, and missing responses were not modeled. We also included nuisance predictors of 6 motion correction parameters (translation and rotation in the x, y, and z directions) in the GLM to account for influence of head motions on the neural activity. In a second GLM, we modeled the neural response to the variation of trial-wise subjective value of the lottery by including the subjective value as a parametric modulator for each of the four decision-condition binary predictors. Subjective value of the lottery in each trial was calculated uniquely for each participant by equation (1), by taking the fitted α and β for each participant under each domain of either gains or losses. Because we fitted the choice data in the loss domain by inputting the positive outcome value, we flipped the sign of the calculated subjective value back in the loss domain. We calculated the subjective values taking α’s and β’s fitted from the two sessions separately, because it would make the estimate of neural response to subjective value variation more accurate. Subjective values were normalized within each scanning block before GLM fitting, so that the estimated effect reflected each participant’s neural response to the variation of subjective value, rather than to its absolute magnitude. Predictor of choice response and nuisance predictors of motion correction were included as in the first GLM. In the third and fourth GLMs, we aimed to further investigate the shape of the neural representation of subjective values. In both GLMs, we combined trials of gains and losses, and only separated trials by uncertainty types. Thus, we included two binary predictors, ambiguous trials and risky trials, in both GLMs, and modeled them as box-car functions with a duration of choice display (6TR). In the third GLM, we included the subjective value itself as a parametric modulator to accompany each binary predictor, to look at the monotonic value-encoding of subjective values. In the fourth GLM, we included the absolute value of subjective value as a parametric modulator to accompany each binary predictor, to look at the U-shaped saliency-encoding of subjective values. The predictor of choice response and nuisance predictors of motion correction were included as in the first GLM. In the fifth GLM to more directly visualize the subjective-value encoding pattern, we made binary predictors based on the subjective value of the lottery. For each participant, we first separated all trials into risky and ambiguous one. Within each uncertainty domain, we then grouped loss trials into 3 bins, by comparing the subjective value of the lottery in each trial to the 1/3 and 2/3 quantile value of the subjective values of all the loss lotteries in this uncertainty domain. Similarly, we grouped gain trials into 3 bins, by comparing the subjective value of the lottery in each trial to the 1/3 and 2/3 quantile value of the subjective values of all the gain lotteries in this uncertainty domain. We then constructed a binary predictor for each bin as a box-car function with the duration of choice display (6TR). Altogether this GLM included 12 predictors (2 uncertainty domains × 2 gain/loss domain × 3 bins) representing the levels of subjective values. Within each uncertainty domain, there were 6 bins of trials, and the 1^st^ bin included the loss lotteries with the most negative subjective values, and the 6^th^ bin included the gain lotteries with the most positive subjective values. An additional predictor of response was modeled as the same way as the other GLMs.

In the second-level analysis, random-effect group analysis was conducted to test whether the mean effect of interest was significantly different from zero across participants, or significantly different between groups by contrasting veterans with PTSD and combat controls. We also took a dimensional approach to test whether the predictor effects were related to the severity of PTSD and other clinical symptoms. The tests were conducted both in a whole-brain search and in ROIs. All whole-brain statistical maps were thresholded at p < 0.001 per voxel, and corrected for multiple comparisons using cluster-extent correction methods through Alphasim by AFNI (58) to control family-wise error (FWE) rate at 0.05. After identifying regions from the whole-brain analysis, in which the neural representation of subjective values was influenced by PTSD symptom severity, we took a leave-one-subject-out (LOSO) approach to define these ROIs in an un-biased way for each participant. For each left-out participant, we defined an ROI from a whole-brain analysis using data from all other participants, so this ROI definition was not influenced by the left-out participant. We then sampled neural signals of the left-out participant’s data from this ROI. We repeated the process for all participants.

## Acknowledgements

This research project is supported by National Center for PTSD, and NIMH Grant R21MH102634. We also thank graduate student training grants from the Interdepartmental Neuroscience Program at Yale and the China Scholarship Council. We thank Alicia Roy and Dr. Erin O’Brien for the recruitment and clinical assessment of veterans; Dr. Michael A. Grubb for development of the task paradigm; Simon Podhajsky, Yumiko Nakamura, and Pooja Salhotra for help with running experiments and data pre-processing. We also thank Dr. Jutta Joormann and Dr. Hyojung Seo for suggestions and comments on data analyses.

