## Supplementary information for "From Value to Saliency: Neural Computations of Subjective Value under Uncertainty in PTSD"

### Supplementary Methods

#### Participants data exclusion

68 male veterans (ages: 23.6-74.6; mean ± standard deviation: 39.4 ± 11.5), who had been deployed and exposed to combat, were recruited. Participants either had current diagnoses of PTSD at the time of the study or were never diagnosed with PTSD (controls). Due to the small proportion of female combat veterans (15% of female in Army 2019, statistics from Department of Defense), we only included male participants. PTSD diagnosis was based on the Clinician Administered PTSD Scale for DSM-4 (CAPS) (1). Data from 10 participants were excluded due to a large number of missing responses or low correct response rate in catch trials in the main task (see Manipulation check below), resulting in 58 participants (ages: 23.6-67.0; amean ± standard deviation: 37.3 ± 8.9) whose behavioral data were reported. Among these 58 participants, CAPS was missing for one PTSD participant, so we only included him in the group comparison but not the dimensional analysis. Imaging data from 10 additional participants were excluded due to excessive movement in the scanner, or because their data were collected using different scanning parameters (5 participants), resulting in 48 participants (ages: 23.6-67.0; mean ± standard deviation: 37.4 ± 9.2), whose neural results were reported. Full characteristics of the sample included in the analysis were reported in Table S1 (behavioral) and Table S2 (neural). Seven participants also took part in a previous behavioral study (2) using a similar paradigm, and these participants were included in both behavioral and imaging analyses. Two of these participants were diagnosed as PTSD in the previous behavioral study but were grouped into combat controls who never developed PTSD in this imaging study. This discrepancy could be due to the inaccuracy of subjective report at the time of the behavioral study, or of the imaging study, and could not be resolved. Behavioral results excluding these seven recurring participants are also reported in Fig S3.

For clinical assessment, we did not exclude veterans with history of substance abuse, to allow for a diverse representative sample of trauma-exposed symptomatology. However, we conducted urine test and breathalyzer for anyone with substance abuse history, or if we suspected any intoxication, and excluded those with positive results.

#### Decision making under risk and ambiguity

##### Task design

The experimental design was based on a previous neuroimaging study (3) and similar to the design of a previous behavioral study in combat veterans (2).

The experiment consisted of choices about risky and ambiguous gains and losses. On gain trials, participants made choices between a fixed monetary gain ($5) and a lottery with chance of a monetary gain but also chance of no gain ($0) (Fig 1B, left). Lottery outcome probability was represented by an image of a rectangle with blue and red areas. On each trial, one color was associated with a monetary gain, and the other color was associated with the null outcome ($0). The size of each colored area represented the probability of getting the outcome associated with it. In half of the trials, outcome probability was fully known (25%, 50% or 75%; 'Risk', Fig 1C left). In the other half of the trials, probability was only partially known. This was achieved by covering the middle part of the colored image with a grey bar ('Ambiguity', Fig 1C right). On different trials the bar covered 24%, 50% or 74% of the image, creating three levels of ambiguity. For example, the lower left example in Fig 1B represents a lottery with a chance between 25% and 75% of winning $27. Each of the risk and ambiguity levels corresponded to an actual physical bag with a total number of 100 red and blue chips inside. For risky bags (corresponding to the risk level images), the exact numbers of red and blue chips were equal to the numbers shown on the images. For ambiguous bags (corresponding to the ambiguity level images), the exact numbers of red and blue chips were within the range shown on the image. Participants were shown these bags during study introduction and were informed that the lottery images they saw during the task corresponded to these bags. The bags would be used to actualize participants' choice, and they were free to inspect the contents of the bags after finishing the study. The potential monetary gain of the lottery varied within a wide range ($5, 6, 7, 8, 10, 12, 14, 16, 19, 23, 27,

31, 37, 44, 52, 61, 73, 86, 101, and 120). The non-zero monetary outcome was randomly related to red or blue, so color was independent from outcome preference.

Loss trials were similar to gain trials, except that participants chose between losing \$5 for sure, and playing a lottery with chance of losing money, but also chance of not losing any money (\$0) (Fig 1B, right). Same as gain trials, half of the loss lotteries were risky and half of them were ambiguous, and the outcome probability was presented in the same way. The potential monetary loss of the lottery varied within the same range but of negative monetary outcomes (–\$5, 6, 7, 8, 10, 12, 14, 16, 19, 23, 27, 31, 37, 44, 52, 61, 73, 86, 101, and 120).

Fig 1D illustrates the timing of each trial. Participants had 6 seconds to look at the options and consider their decision. A green circle then appeared in the middle of the screen, cueing participants to make a choice within 3.5 seconds. If they did not register a response, the response would be counted as missing. A feedback image of two side-by-side squares was shown for 0.5 second immediately following the button press to confirm the choice, with the yellow square indicating the side participants chose. The feedback was followed by a jittered inter-trial-interval (ITI) of a white circle, the duration of which was among 4, 6, 8 seconds plus the remainder of the response time limit (3.5s – reaction time (RT)). To avoid learning, chosen lotteries were not played during the scan. In both the gain and the loss domains, all risk and ambiguity levels were paired with all amount levels, resulting in 120 unique gain trials ((3 + 3) × 20) and 120 unique loss trials. Each trial type was presented once. All trials were grouped into 8 blocks of 30 trials each. Each block contained either only gain trials or loss trials, but risky and ambiguous trials were mixed in a random order. This resulted in 4 gain blocks and 4 loss blocks. Due to the length of the experiment, these blocks were divided into two scanning sessions on two separate days. The interval between the two scanning sessions was on average 12.6 days.

#### Manipulation check

To verify that participants understood the task, and that they aimed to maximize earnings and minimize losses, we included 12 trials in which the potential lottery outcome was identical to the certain amount (±$5). In these trials, one option is clearly better than the other (e.g. a certain gain of $5 should be preferred over a 50% chance of gaining $5). Data from participants who chose the inferior option on more than 50% of these trials were excluded.

#### Task administration

On Day1, participants were introduced to the task, and were required to correctly respond to several questions to make sure they understand the task and the lottery presentation. They also practiced 16 trials before the actual task. After the introduction, each participant was endowed with $120 (the maximal possible loss), and then went through two gain and two loss blocks, whose order was counterbalanced across participants (either Gain-Gain-Loss-Loss or Loss-Loss-Gain-Gain). Each block contained 30 trials together with an additional trial in the beginning (a choice between +/-$5 and a lottery offering +/-$4) to capture the initial burst of activity. The added one trial was excluded from analysis.

On Day2, participants were first reminded of the task, and then went through another four blocks of choices (two gain and two loss), in an opposite order to what they had on Day1. Following the scanning, one trial out of the 240 trials (including both gains and losses) was randomly selected and realized. If the participant chose the sure option, $5 were added to or subtracted from the $120 endowment. If the participant chose the lottery, he\she would play the lottery by pulling a chip out of the physical bag corresponding to the lottery image, and the outcome related to the color of the pulled-out chip would be added to or subtracted from the $120. At the end of the second session, KBIT, demographic questionnaires, BIS/BAS, and BIS-11 were collected.

#### Analysis

##### Analysis of clinical symptoms and trauma-related measures

We used the 5-factor model of PTSD (4) to assess multidimensional PTSD symptoms, including re-experiencing, avoidance, emotional numbing, dysphoric arousal, and anxious arousal. We calculated both the overall symptom severity, and the 5 factors' symptom severities by breaking down the questionnaire items. To account for comorbidities, we conducted principal component analysis (PCA) on all clinical and trauma-exposure measurements, including the 5 factors of CAPS, Beck's Depression Inventory (BDI), state and trait anxiety indexes separately from State-Trait Anxiety Inventory (STAI), Dissociative Experiences Scale (DES), Combat Exposure Scale (CES), and Childhood Trauma Questionnaire (CTQ).

##### Model-based risk and ambiguity attitudes analysis

The model-fitting was conducted in MATLAB (Version R2018b) through maximum likelihood. We primarily used MATLAB function fminunc to minimize the negative log-likelihood function, and switched to using fminsearch if it failed to converge. Under our task design, we could detect risk parameters ($\alpha$) in the range of [0.0905, 7.6036], and ambiguity parameters $\beta$ in the range of [-4.0303, 4.1667]. These constraints were calculated based on the range of uncertainty levels and monetary magnitudes in our task design, same as the procedure used in previous studies (2). The lower boundary of $\alpha$ was determined by equating the subjective value of the best lottery (75% chance of \$120) with the subjective value of the certain option (\$5), using equation (1). Similarly, to determine the upper boundary of $\alpha$, we equated the subjective values of the worst lottery (25% chance of \$6) and the certain option (\$5). Because the choices of ambiguous lotteries also depend on the risk attitude $\alpha$, the boundary values of $\beta$ also depend on $\alpha$. So we simply calculated all possible $\beta$'s by equating the subjective values of each ambiguous lottery and the certain option (\$5), using the boundary values of $\alpha$. The boundary values of $\beta$ were then determined by the minimum and maximum of all calculated $\beta$'s. Even without constraining $\alpha$ and $\beta$ during the modeling fitting procedure, the fitted results of all participants included in behavioral and imaging analyses were within this range. We fitted each participant's choices combining data from two sessions and obtained four attitudes:

risk attitudes for gains and losses, ambiguity attitudes for gains and losses. For consistency, we transformed all attitudes in the following way such that negative values indicate aversion and positive values indicate seeking: risky gains: $\alpha - 1$, risky losses: $1 - \alpha$, ambiguous gains: $-\beta$, ambiguous losses: $\beta$. Since participants performed the task on two separate sessions, we also fitted each session's choice data separately. These fitted parameters from separate sessions were used to calculate trial-wise subjective values of the lotteries for GLM neural analysis, because they could capture the subjective values more accurately for searching neural activity change induced by variations of subjective values.

We then analyzed the fitted risk and ambiguity attitudes both in group comparisons between veterans with PTSD and combat controls, and through a dimensional approach by looking at their correlation with clinical symptoms and other continuous measurements. We also accounted for the potential influence of demographic factors (age, income, education, and intelligence) on uncertainty attitudes, by conducting multi-factor ANOVAs through a Generalized Linear Model:

Uncertainty attitude in one decision condition (e.g. Risk attitude in gains) ~ CAPS total + age + income (categorical) + education (categorical) + intelligence + BIC

In the data we collected, age and intelligence were continuous variables, income was a categorical variable with 10 possible levels, and education was a categorical variable with 6 possible levels. All continuous variables were standardized before fitting the linear model.

#### fMRI GLM first-level analysis

Analysis of fMRI data were conducted in the Neuroelf toolbox (Version 1.0, https://neuroelf.net/) through MATLAB (Version R2018b) for the purpose of fitting generalized linear models (GLM), extracting regions-of-interest (ROI) data, and visualizing brain statistical maps. Further statistical analyses and visualization were conducted in R (Version 3.5.1) (5) with packages ez (6), psych (7), nlme (8), emmeans (9), ggplot2 (10), and PerformanceAnalytics (11). We investigated the neural response under different decision conditions by fitting pre-processed functional signals with generalized linear models (GLM). The pre-processed fMRI signal time course was first converted to percent signal change within each scanning block, and activity of each voxel was modeled by GLM predictors convolved with a standard double-gamma hemodynamic response function.

In the first GLM, we looked at the general activity during decision making, by including four binary predictors for all four decision conditions: ambiguous gains, risky gains, ambiguous losses, and risky losses. Each binary predictor was modeled as a box-car function, with the duration of choice display (6TR). We modeled choice response of all trials by another binary predictor with the duration of 1TR at the time of button press, and missing responses were not modeled. We also included nuisance predictors of 6 motion correction parameters (translation and rotation in the x, y, and z directions) in the GLM to account for influence of head motions on the neural activity.

In a second GLM, we modeled the neural response to the variation of trial-wise subjective value of the lottery by including the subjective value as a parametric modulator for each of the four decision-condition binary predictors. Subjective value of the lottery in each trial was calculated uniquely for each participant by equation (1) (See section Model-based risk and ambiguity attitudes estimation), by taking the fitted $\alpha$ and $\beta$ for each participant under each domain of either gains or losses. Because we fitted the choice data in the loss domain by inputting the positive outcome value, we flipped the sign of the calculated subjective value back in the loss domain. We calculated the subjective values taking $\alpha$'s and $\beta$'s fitted from the two sessions separately, because it would make the estimate of neural response to subjective value variation more accurate. Subjective values were normalized within each scanning block before GLM fitting, so that the estimated effect reflected each participant's neural response to the variation of subjective value, rather than to its absolute magnitude. Predictor of choice response and nuisance predictors of motion correction were included as in the first GLM.

In the third and fourth GLMs, we aimed to further investigate the shape of the neural representation of subjective values. In both GLMs, we compared gains and losses on the same scale, and only separated trials by uncertainty types. Thus, we included two binary predictors, ambiguous trials and risky trials, in both GLMs, and modeled them as box-car functions with a duration of choice display (6TR). In the third GLM, we included the subjective value itself as a parametric modulator to accompany each binary predictor, to look at the monotonic value-encoding of subjective values. In the fourth GLM, we included the absolute value of subjective value as a parametric modulator to accompany each binary predictor, to look at the U-shaped saliency-encoding of subjective values. The predictor of choice response and nuisance predictors of motion correction were included as in the first GLM.

In the fifth GLM, we aimed to more directly visualize the subjective-value encoding pattern. We made binary predictors based on the subjective values of the lottery. For each participant, we first separated all trials into risky and ambiguous trials. Within each uncertainty domain, we then grouped loss trials into 3 bins, by comparing the subjective value of the lottery in each trial to the 1/3 and 2/3 quantile value of the subjective values of all the loss lotteries in this uncertainty domain. Similarly, we grouped gain trials into 3 bins, by comparing the subjective value of the lottery each trial to the 1/3 and 2/3 quantile value of the subjective values of all the gain lotteries in this uncertainty domain. We then constructed a binary predictor for each bin as a box-car function with the duration of choice display (6TR). Altogether this GLM included 12 predictors (2 uncertainty domains × 2 gain/loss domain × 3 bins) representing the levels of subjective values. Within each uncertainty domain, there were 6 bins of trials, and the 1st bin included the loss lotteries with the most negative subjective values, and the 6th bin included the gain lotteries with the most positive subjective values. An additional predictor of response was modeled in the same way as the other GLMs.

#### fMRI GLM second-level analysis

After individual GLM fitting, random-effect group analysis was conducted to test whether the mean effect of interest was significantly different from zero across participants, or significantly different between groups, by contrasting veterans with PTSD and combat controls. We also took a dimensional approach to test whether the predictor effects were related to the severity of PTSD and other clinical symptoms.. The tests were conducted both in a whole-brain search and in ROIs.

In whole-brain analyses, all statistical maps were thresholded at $p < 0.001$ per voxel, and corrected for multiple comparisons using cluster-extent correction methods through Alphasim by AFNI (12) to control family-wise error (FWE) rate at 0.05.

In ROI analyses, investigating the general neural activity during decision making based on the first GLM, we chose the region in vmPFC whose activity showed negative correlation with CAPS total in the whole-brain analysis. We further investigated which factor of the PTSD symptoms drove this negative relationship, by fitting a linear model including all five symptoms based on the CAPS measurements together with age and intelligence:

Averaged GLM beta over all four decision conditions ~ re-experiencing + avoidance + emotional numbing + dysphoric arousal + anxious arousal + age + intelligence

We then conducted variables subset selection to identify which symptom cluster(s) best influenced vmPFC neural activity, using exhaustive search through the package “leaps” (13) in R. We compared regression models including all possible combinations of variables for each given number of predictors of this linear model (ranging from including only one predictor to including all seven predictors), and selected the best model with the lowest Bayesian Information Criterion (BIC). The best linear model identified by this exhaustive approach could identify both the best number of symptom clusters to include, as well as which symptom cluster(s).

In ROI analyses, investigating subjective value representations based on the second, third and fourth GLMs, we chose two brain regions, vmPFC and ventral striatum, based on the meta-analysis by Bartra et al. (14), which identified these areas as encoding the value of different reward categories. To control for the potential influence of demographic factors on neural activity, we conducted multi-factor ANOVAs through a Generalized Linear Model that included the PTSD total symptom severity and demographic factors, to explain the neural subjective value signal, similar to the behavioral analysis of uncertainty attitudes:

Neural subjective value signal of a type of lottery (e.g. risk gain lottery) ~ CAPS total + age + income (categorical) + education (categorical) + intelligence

All continuous variables were standardized before fitting the linear models.

#### Leave-one-subject-out (LOSO) procedure

After identifying regions from the whole-brain analysis, in which the neural representation of subjective values was influenced by PTSD symptom severity, we took a leave-one-subject-out (LOSO) approach to define these ROIs in an un-biased way for each participant. For each left-out participant, we defined an ROI from a whole-brain analysis using data from all other participants, so this ROI definition was not influenced by the left-out participant. We then sampled neural signals of the left-out participant's data from this ROI. We repeated the process for all participants.

#### Using behavioral and neural measures to predict PTSD symptom variation

To investigate whether symptom variation could be better explained by behavioral uncertainty attitudes or by neural activity during decision making, we constructed an additional model. This model included the betas of each decision condition (ambiguous losses, risky losses, ambiguous gains, and risky gains) in the first GLM, sampled from the vmPFC area identified in the whole-brain analysis:

Neural model: CAPS total ~ neural activity under risky gains + neural activity under ambiguous gains + neural activity under risky losses+ neural activity under ambiguous losses + age + intelligence

Similarly, we constructed a linear model using behavioral uncertainty attitudes in the four decision conditions to explain PTSD symptom severity:

Behavioral model: CAPS total ~ risk attitude in gains + ambiguity attitude in gains + risk attitude in losses + ambiguity attitude in losses + age + intelligence

We also constructed a full model including both neural and behavioral measures:

Full model: CAPS total ~ risk attitude in gains + ambiguity attitude in gains + risk attitude in losses + ambiguity attitude in losses + neural activity under risky gains + neural activity under ambiguous gains + neural activity under risky losses+ neural activity under ambiguous losses + age + intelligence

All variables were standardized before fitting the linear model. All models included age and intelligence to control for these two demographic factors. We compared these three models by BIC.

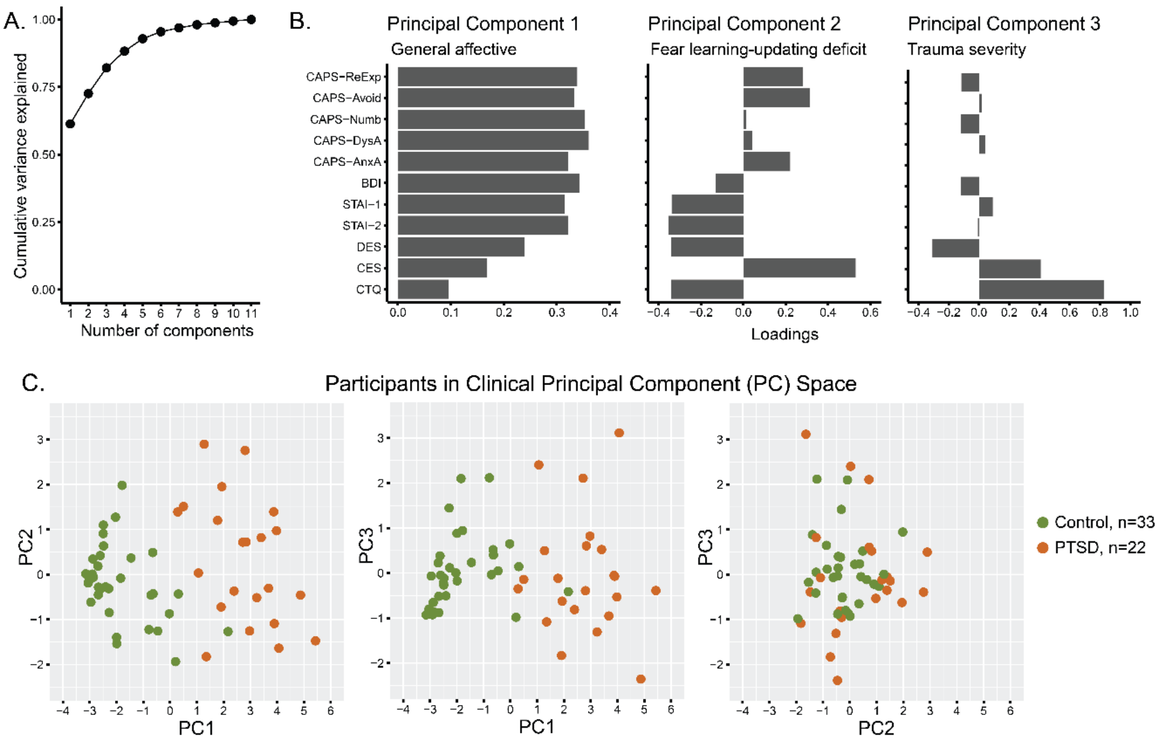


**Fig. S1.** Principal component analysis (PCA) of clinical symptoms and trauma exposure measures. A: Cumulative variance explained by all principal components. B: Loading coefficients of the first three principal components, representing general affective symptoms, deficit in fear learning-updating, and trauma severity, respectively. Labels: CAPS-ReExp: re-experiencing, CAPS-Avoid: avoidance, CAPS-Numb: emotional numbing, CAPS-DysA: dysphoric arousal, CAPS-AnxA: anxious arousal, BDI: Beck Depression Inventory, STAI-1: State Anxiety, STAI-2: Trait Anxiety, DES: Dissociative Experiences Scale, CES: Combat Exposure Scale, CTQ: Childhood Trauma Questionnaire. C: Participants plotted in the two-dimensional spaces represented by pairs of the first three principal components, colored by PTSD diagnosis.

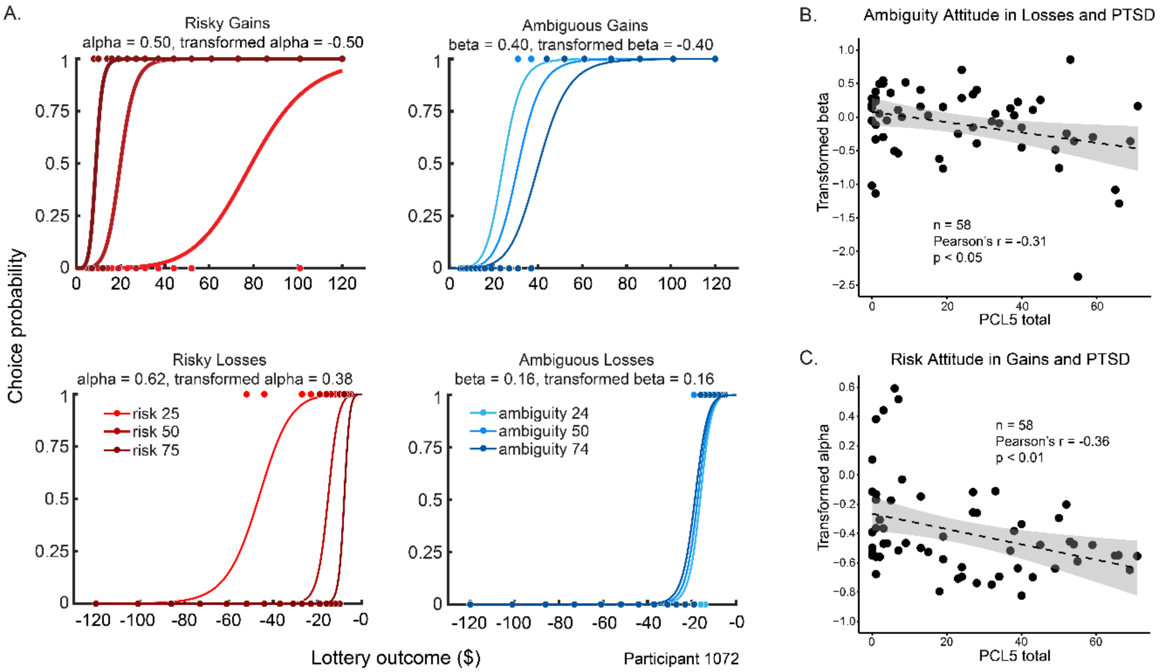


**Fig. S2.** Additional behavioral results. A: An example of behavioral model fitting from a single participant. Dots on the graphs represent the actual choices (1 = choosing the lottery, 0 = choosing the certain option) of each type of lottery. Color shade represents the level of risk (red) and ambiguity (blue). Curves are the fitted choice probability as a function of lottery outcome. The sign of transformed attitudes indicates seeking (positive) or aversion (negative) for both risk and ambiguity. This participant was risk averse in gains, risk seeking in losses, ambiguity averse in gains, and ambiguity seeking in losses. B: Ambiguity attitude in losses was negatively correlated with PTSD symptom severity indicated by PCL5. C: Risk attitude in gains was negatively correlated with PTSD symptom severity indicated by PCL5.

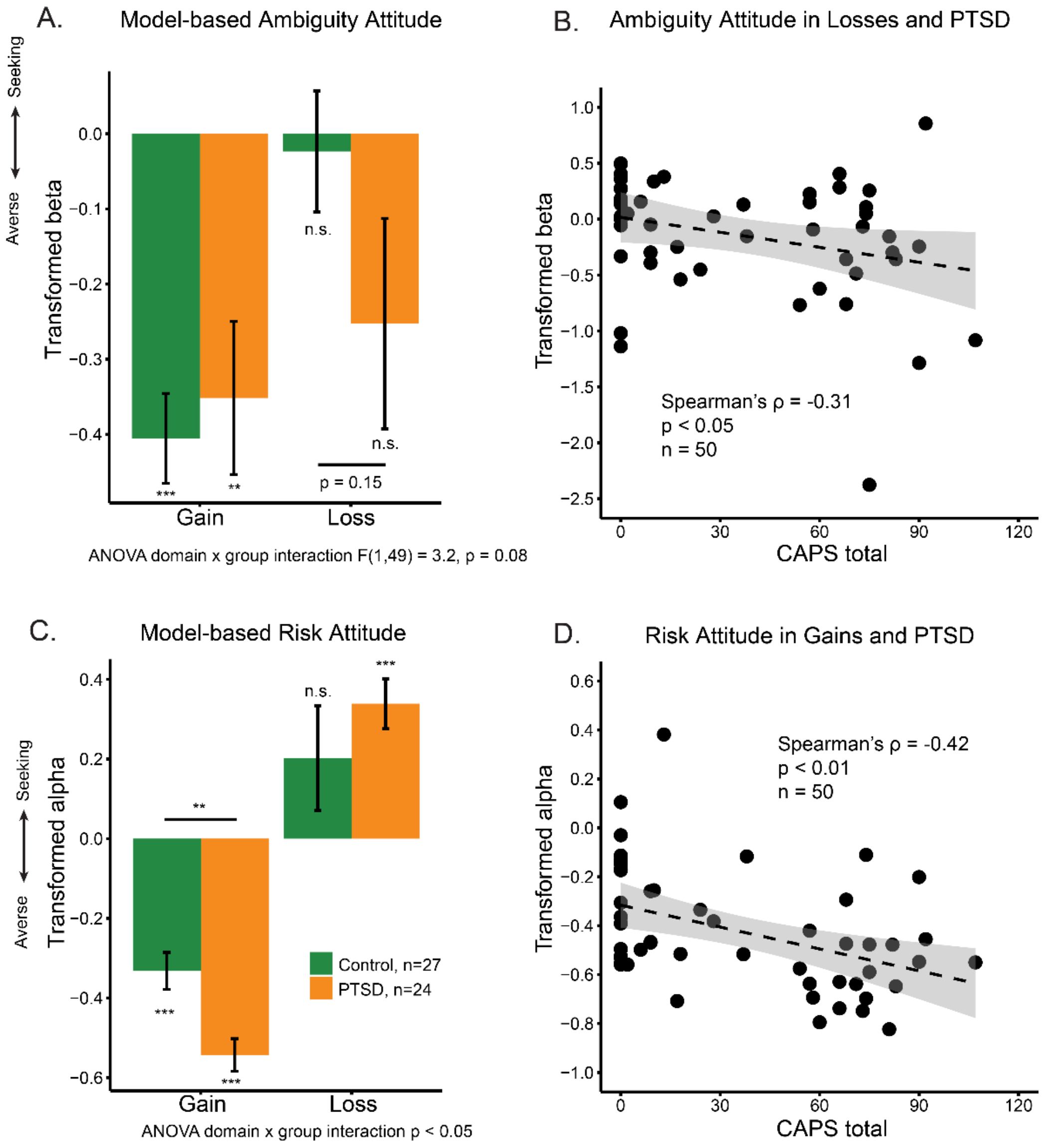


**Fig. S3.** Uncertainty attitudes and PTSD symptom severity, excluding 7 combat control veterans who also participated in the previous behavioral study (2). A: Group comparison between veterans with PTSD and combat controls of ambiguity attitudes in gains and losses. B: PTSD symptom severity was negatively correlated with ambiguity attitude in losses. C: Group comparison between veterans with PTSD and combat controls of risk attitudes in gains and losses. D: PTSD symptom severity was negatively correlated with risk attitude in gains. In A and C, comparing each group's attitudes with zero were FDR-corrected across all four comparisons in each uncertainty type. Post-hoc comparisons between groups were FDR-corrected after conducting ANOVA. Significance level: *, $p<0.05$; **, $p<0.01$; ***, $p<0.001$.

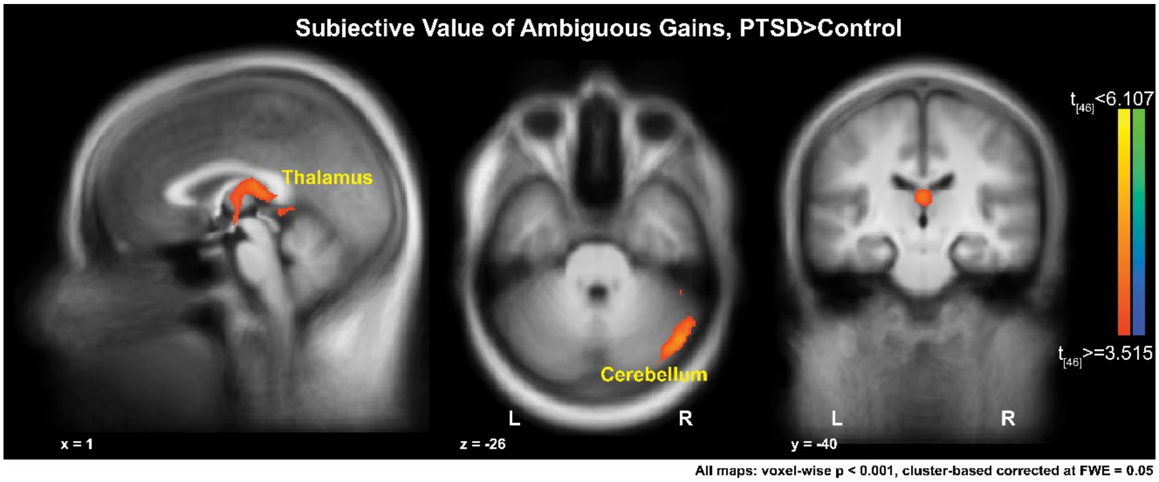


**Fig. S4.** A whole-brain comparison of neural subjective-value signals of ambiguous gains between veterans with PTSD and combat controls by direct contrast. The map was corrected using cluster-based method controlling for family-wise error at 0.05, when thresholded at $p < 0.001$ at the voxel level.

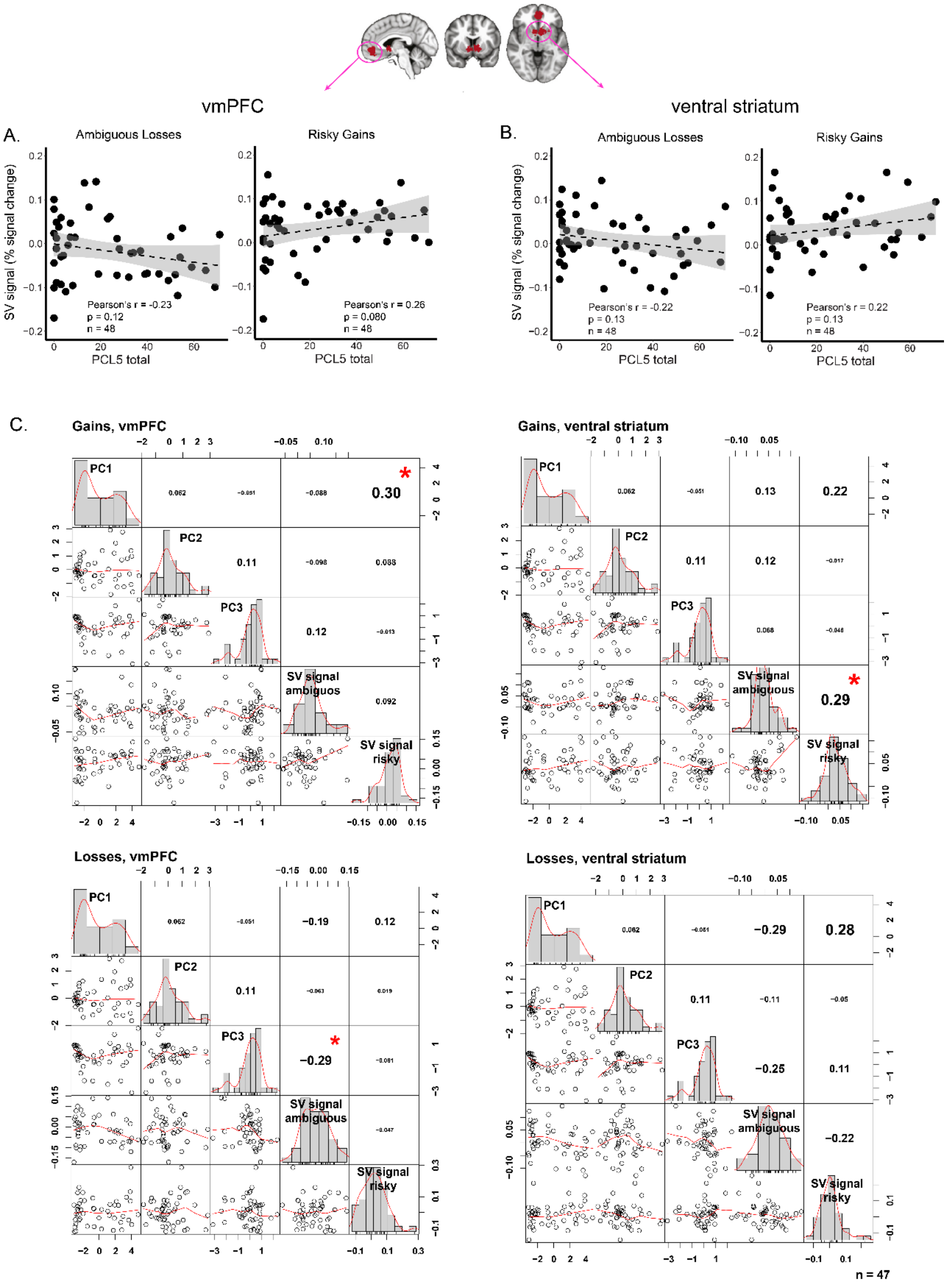


**Fig. S5.** Additional analyses of subjective-value representation in external ROIs of vmPFC and ventral striatum. A: In vmPFC, correlations between subjective-value representation of ambiguous losses and risky gains separately with PTSD symptom severity indicated by PCL5. B: In ventral striatum, correlations between subjective-value representation of ambiguous losses and risky gains separately with PTSD symptom severity indicated by PCL5.  C: Correlations between subjective-value neural representation and PCA components from all clinical and trauma measures. Figures were organized by domain (gains top, losses bottom) and ROIs (vmPFC left, ventral striatum right). Each cell of the graph is a pairwise correlation between the variables. Numbers in the upper right cells indicate Pearson's correlation. Significance level: *, $p<0.05$.

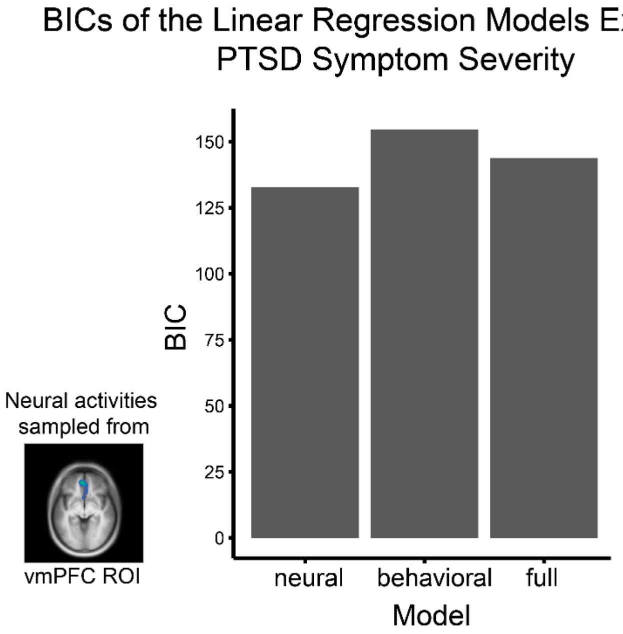


**Fig. S6.** Variation of PTSD symptom severity was better explained by neural activities than behavioral attitudes. Comparing BICs of linear models including (1) only general neural activities under four decision conditions in vmPFC area defined in Fig 4, (2) only behavioral uncertainty attitudes under four decision conditions, (3) both general neural activities and behavioral uncertainty attitudes under four decision conditions, to explain PTSD symptom severity indicated by CAPS total score. The model including only the neural measures performed the best.

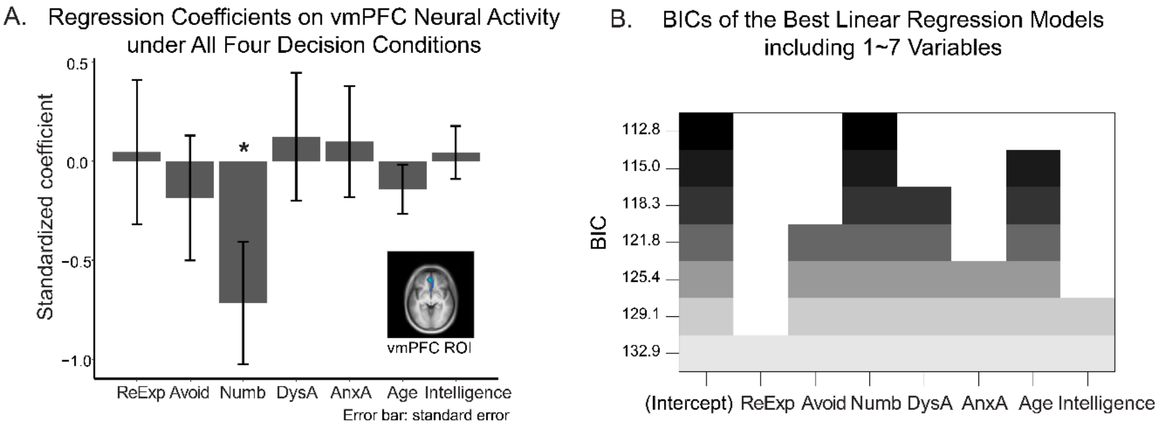


**Fig. S7.** Emotional numbing plays the key role in diminished vmPFC activity. A: Emotional numbing symptom severity drove the negative relationship between vmPFC neural activity and PTSD symptom severity (Fig 4), revealed by a linear regression model on the vmPFC activity including all clusters of the 5-factor model of CAPS. B: Variable selection using exhaustive search also indicated that emotional numbing was the key symptom driving this relationship. Each row of the graph shows the selected variables (shaded) for the best model with a given number of predictors. Rows are ranked and colored by BIC. The top row represents the best model, which includes only Emotional numbing as the predictor, among all possible combinations of predictors. Variable naming: ReExp: re-experiencing, Avoid: avoidance, Numb: emotional numbing, DysA: dysphoric arousal, AnxA: anxious arousal.

**Table S1.** Descriptive statistics of demographics and clinical measures of participants reported in the behavioral results.

| | **PTSD** | **Control** |
|---|---|---|
| Number of participants | 24 | 34 |
| Age | 34.70 (6.44) | 39.17 (10.03) |
| Kaufman Brief Intelligence Test (KBIT) | 105.63 (10.52) | 111.59 (13.22) |
| Clinician Administered PTSD Scale (CAPS)-Total Score | 72.13 (15.04) | 6.21 (9.68) |
| CAPS-Re-experiencing | 19.83 (6.79), n = 23 | 1.00 (1.86) |
| CAPS-Avoidance | 10.65 (3.41), n = 23 | 0.47 (1.38) |
| CAPS-Emotional Numbing | 18.13 (6.61), n = 23 | 1.03 (3.03) |
| CAPS-Dysphoric Arousal | 15.09 (3.15), n = 23 | 2.00 (3.65) |
| CAPS-Anxious Arousal | 8.44 (2.56), n = 23 | 1.71 (2.87) |
| PTSD Checklist for DSM-5 (PCL-5) | 41.54 (15.79) | 10.88 (15.95) |
| Beck Depression Inventory (BDI) | 26.11 (13.80) | 5.50 (7.88) |
| State Anxiety (STAI-1) | 47.92 (12.72) | 32.62 (9.43) |
| Trait Anxiety (STAI-2) | 47.32 (16.23) | 31.33 (10.94) |
| Dissociative Experience Scale (DES) | 44.04 (34.53) | 17.31 (20.75), n = 33 |
| Combat Exposure Scale (CES) | 19.96 (9.06), n = 23 | 13.00 (8.57) |
| Childhood Trauma Questionnaire (CTQ) | 39.04 (14.03), n = 23 | 34.43 (8.76) |

Mean (standard deviation)

Fewer number of participants for several measures was due to incomplete data.

**Table S2.** Descriptive statistics of demographics and clinical measures of participants reported in the neural results.

| | **PTSD** | **Control** |
|---|---|---|
| Number of participants | 19 | 28 |
| Age | 35.59 (6.86) | 38.62 (10.42) |
| Kaufman Brief Intelligence Test (KBIT) | 104.16 (11.24) | 112.97 (13.42) |
| Clinician Administered PTSD Scale (CAPS)-Total Score | 71.42 (15.52) | 5.55 (9.40) |
| CAPS-Re-experiencing | 19.84 (7.41) | 1.00 (1.95) |
| CAPS-Avoidance | 10.21 (3.52) | 0.38 (1.21) |
| CAPS-Emotional Numbing | 17.95 (6.91) | 0.72 (2.14) |
| CAPS-Dysphoric Arousal | 14.89 (2.81) | 1.79 (3.40) |
| CAPS-Anxious Arousal | 8.53 (2.41) | 1.66 (2.93) |
| PTSD Checklist for DSM-5 (PCL-5) | 42.37 (15.46) | 9.93 (15.99) |
| Beck Depression Inventory (BDI) | 24.95 (13.40) | 5.07 (8.08) |
| State Anxiety (STAI-1) | 46.96 (11.41) | 31.69 (8.69) |
| Trait Anxiety (STAI-2) | 45.62 (16.07) | 30.12 (10.32) |
| Dissociative Experience Scale (DES) | 47.53 (36.50) | 16.17 (20.61) |
| Combat Exposure Scale (CES) | 21.06 (9.70), n = 18 | 13.14 (8.19) |
| Childhood Trauma Questionnaire (CTQ) | 39.05 (15.53), n = 18 | 34.64 (9.25) |

Mean (standard deviation)

For imaging analysis, 10 participants were excluded from those reported in the behavioral results because of fMRI data quality. Fewer number of participants for several measures was due to incomplete data.

**Table S3.** All areas showing whole-brain contrast of subjective value representation, comparing veterans with PTSD (n = 19) and combat controls (n = 29).

| Condition | Contrast | Region | Side | Mean t statistic | Peak Talairach coordinates | | | Cluster size |
|---|---|---|---|---|---|---|---|---|
| | | | | | x | y | z | |
| Risky gain | PTSD > Control | Orbital frontal cortex | R | 4.06 | 31 | 55 | 0 | 321 |
| Ambiguous gain | PTSD > Control | Cerebellum | R | 3.91 | 47 | -65 | -26 | 238 |
| | | Thalamus | L/R | 3.78 | -1 | -21 | 14 | 396 |
| Risky Loss | PTSD > Control | None | N/A | N/A | N/A | N/A | N/A | N/A |
| Ambiguous Loss | Control > PTSD | Cuneus | R | 3.91 | 15 | -99 | 4 | 537 |
| | | Inferior frontal gyrus | L | 3.79 | -53 | 19 | -2 | 197 |
| | | Middle occipital gyrus | L | 3.75 | -33 | -91 | 6 | 184 |

Areas were defined by statistical maps corrected using cluster-based method controlling family-wise error at 0.05 when thresholded by $p < 0.001$ at the voxel level.
